# Discovery and characterization of a lactonase in gut microbiota that initiates the metabolism of ellagic acid

**DOI:** 10.64898/2025.12.05.692447

**Authors:** Zhike Xie, Jiaxin You, Feifei Xin, Feng Chen, Qiangqiang Ma, Yiping Guo, Zheng Ruan

## Abstract

Ellagic acid (EA), a natural plant polyphenol, and its metabolite urolithin (Uro) exhibit significant bioactivity, with Uro being recognized as the active compound responsible for the *in vivo* effects of EA. Although specific intestinal microorganisms that can metabolize EA have been discovered, the corresponding biochemical routes and genetic mechanisms involved in these processes have yet to be fully elucidated. In this study, we employed bioinformatics, biochemical assays, and genetic analyses to discover and characterize an EA lactonase (EAL) from the intestinal bacterium *Gordonibacter urolithinfaciens* DSM 27213. EAL specifically catalyzes the cleavage of the ester bond in EA, yielding Uro-M5. Notably, EAL and its extensive homologs form a unique subclass within the lactonase and amidohydrolase families. Structural predictions indicate that the active pocket of EAL comprises a propeller-like structure formed by seven β-strands. Additionally, molecular modeling analyses further revealed that EA establishes specific interactions with key catalytic residues within the enzyme’s binding site. These insights enhance our understanding of the role of intestinal microbial communities in metabolizing plant-derived phenolic compounds, potentially shaping health outcomes and disease management through dietary interventions.

**Significance Statement:** Ellagic acid (EA), which is abundant in the human gut microbiome, can be metabolized by gut microbes into urolithin A (Uro-A), however, the biochemical mechanisms underlying these transformations remain incompletely understood. In this study, we identified and characterized an EA lactonase (EAL) from the gut bacterium *Gordonibacter urolithinfaciens* DSM 27213, which serves as a key enzyme in the initial metabolic breakdown of EA. We elucidate the function, distribution, and structure of the EAL. This discovery advances our understanding of Uro-A production within the human body and may provide a foundation for further investigations into the relationship between Uros and human health.

## Introduction

The intestinal microbiome significantly contributes to sustaining physiological homeostasis in the host organism through its enzymatic transformation of exogenous compounds, including various phytochemicals like phenolic and flavone compounds, consequently inducing cellular signaling pathways via bacterial metabolites (1, 2). While it is established that the metabolism of these compounds influences health and disease, further mechanistic investigations are needed to fully elucidate the complex enzymatic processes involved in gut microbiota-mediated polyphenol metabolism (3). This need arises from the diversity of polyphenolic components in the diet and the complexity of the gut microbiota (4).

Ellagic acid (EA), a naturally occurring polyphenolic compound primarily found in tannin structures, is widely distributed across various dietary sources, with particularly high concentrations in pomegranates, blackberries, raspberries, and several tree nuts (5, 6). Despite its limited absorption in the human digestive system, EA undergoes significant biotransformation by the gut microbiota, leading to the production of urolithin (Uro) derivatives with notable bioactive potential (7). In the gut, EA undergoes primary bioconversion via lactone ring cleavage, generating the metabolic intermediate Uro-M5. This precursor compound then experiences a sequential series of three distinct hydroxyl group elimination reactions, progressively transforming into Uro-M6, followed by Uro-C, and finally yielding the end product Uro-A (Fig. 2A) (8, 9). Extensive experimental evidence has identified Uro-A as the primary bioactive metabolite responsible for mediating the physiological effects associated with EA (10). The microbial metabolism of EA involves two types of enzymes: lactonases, which facilitate the opening of the lactone ring, and dehydroxylases (11). To date, only Uro-C to Uro-A dehydroxylases have been identified (12).

Current scientific investigations have identified a limited group of microbial species, primarily from the phylum Actinobacteria, that possess the enzymatic capacity to biotransform EA into Uro-C. Among these strains, three well-characterized strains have been extensively studied: *Gordonibacter urolithinfaciens* DSM 27213 (13), along with its phylogenetic relatives *Gordonibacter pamelaeae* DSM 19378 (14) and *Gordoniacter faecis* KGMB12511(15). Additionally, certain potential probiotics, including *Bifidobacterium pseudocatenulatum* INIA P815 (9), *Lactobacillus plantarum* CCFM1286 (16), *Lactococcus garvieae* FUA009 (17), and *Streptococcus thermophilus* FUA329 (18), have demonstrated the ability to transform EA into Uro-A. While the identification of these microbial strains has significantly advanced our comprehension of the physiological significance of EA, the specific enzymes involved in the intestinal microbial catabolism of EA remain incompletely characterized, and the lactonase responsible for the initial step of the metabolic pathway has yet to be elucidated.

In this study, we employed an integrated approach combining bioinformatics and biochemical analyses to identify and characterize an EA lactonase (EAL) derived from *G. urolithinfaciens* DSM 27213. This enzyme has a distinctive substrate specificity, selectively cleaving the cyclic ester bonds of EA. The catalytic mechanism was elucidated through structural prediction of EAL, complemented by molecular docking and site-directed mutagenesis experiments. Importantly, EAL and its widespread homologs constitute a novel class of lactonases/amidohydrolases, which had not been previously recognized.

## Results

### Discovery of EAL in *G*. *urolithinfaciens* DSM 27213

To date, research on EAL and related enzymes remains scarce. Given the sequence similarity among various lactonases, we employed molecular docking techniques to assess the interaction between EA and seventeen types of lactonases from seven different families (Table S1), the docking results indicated that EA was capable of fitting into the active pocket of fourteen of these lactonases (Fig. S1). Successful docking was supported by the docking parameters, as both the -CDOCKER ENERGY and -CDOCKER INTERACTION ENERGY values were greater than 0, while the binding energy values were negative (Table S2). These findings indicate stable and favorable interactions between EA and the active sites of those lactonases. Subsequently, we used the protein sequences of these fourteen lactonases to perform DELTA-BLAST searches against the strain *G. urolithinfaciens* DSM 27213, yielding twenty-seven potential metabolic enzymes (Table S3). We further employed molecular docking to reveal that EA can fit into the active pocket of twenty-five of these potential metabolic enzymes (Fig. 1), and the docking parameters confirmed the reliability of these interactions (Table S4).

**Fig. 1.**
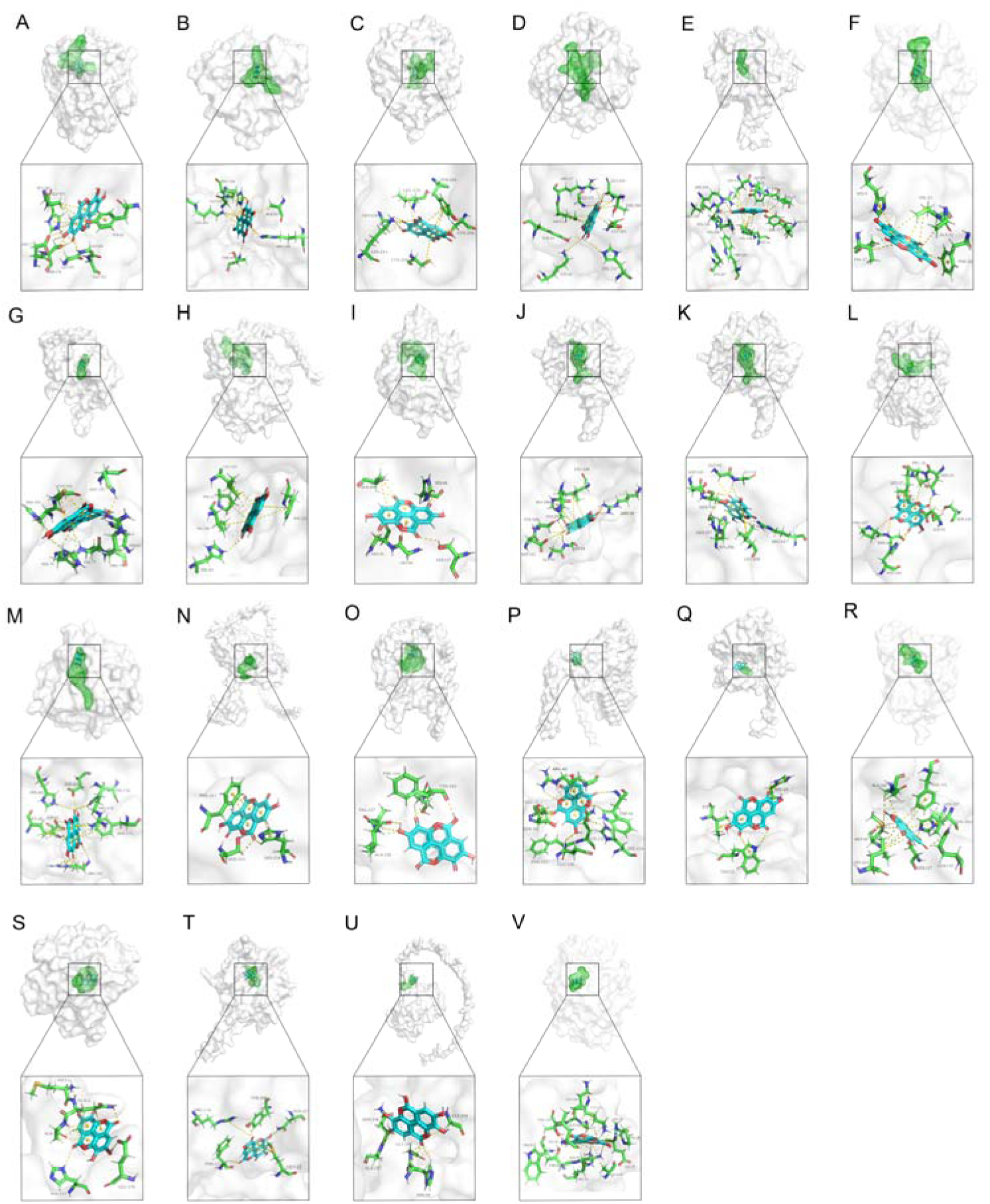
Docking studies of EA with 25 potential EAL identified through DELTA-BLAST in the strain *G. urolithinfaciens* DSM 27213. (A) Acetyl esterase ROT90431.1. (B) Glucosamine-6-phosphate deaminase ROT89736.1. (C) Putative heme d1 biosynthesis radical SAM protein NirJ2 ROT88482.1. (D) 4Fe-4S dicluster domain-containing protein ROT91153.1. (E) Amidohydrolase ROT88165.1. (F) Amidohydrolase ROT91117.1. (G) TatD family deoxyribonuclease ROT89090.1. (H) DUF2974 domain-containing protein ROT92136.1. (I) Alpha/beta hydrolase ROT88025.1. (J) Alpha/beta hydrolase ROT88342.1. (K) Alpha/beta hydrolase ROT88342.1. (L) MBL fold metallo-hydrolase ROT88497.1. (M) MBL fold metallo-hydrolase ROT89054.1. (N) Ribonuclease J ROT88679.1. (O) Lactonase family protein ROT91662.1. (P) Hypothetical protein ROT89966.1. (Q) Hypothetical protein ROT88359.1. (R) Alpha/beta hydrolase ROT90012.1. (S) MBL fold metallo-hydrolase ROT90594.1. (T) Gluconolactonase ROT91236.1. (U) MBL fold metallo-hydrolase ROT89368.1. (V) Peptidase ROT88897.1.

We selected the top six potential metabolic enzymes (ROT90431.1, ROT89736.1, ROT88482.1, ROT91153.1, ROT88165.1, and ROT91117.1) based on binding energy, along with the sole enzyme annotated as a lactonase family protein (ROT91662.1), for further heterologous expression studies (Table S4). These seven target genes (DMP12_05280, DMP12_08305, DMP12_12230, DMP12_03900, DMP12_13210, DMP12_03695, and DMP12_03205) were heterologously expressed in an *Escherichia coli* BL21 (DE3) system to produce the corresponding proteins. After transformation, the recombinant strains were cultured in EA-supplemented lysogeny broth (LB) liquid medium for 24 h, and the metabolic products in the culture supernatant were quantified by high performance liquid chromatography (HPLC) analysis. Notably, the HPLC profile of the supernatant from cultures expressing the DMP12_13210 gene (encoding the ROT88165.1 protein) revealed the appearance of a new peak and consumption of EA, with the new compound identified as Uro-M5 through comparison with the standard (Fig. 2B). The *E. coli* BL21 strain containing this transformant (encoding the ROT88165.1 protein) exhibited a maximum EA conversion rate of approximately 83.86% within 24 h (Fig. 2C and Fig. S2). In contrast, no EA consumption or Uro-M5 formation was observed in the *E. coli* BL21 strains containing any of the other six transformants (Fig. 2D and E). These findings suggested that the DMP12_13210 gene encoded an enzyme responsible for initiating EA catabolism.

**Fig. 2.**
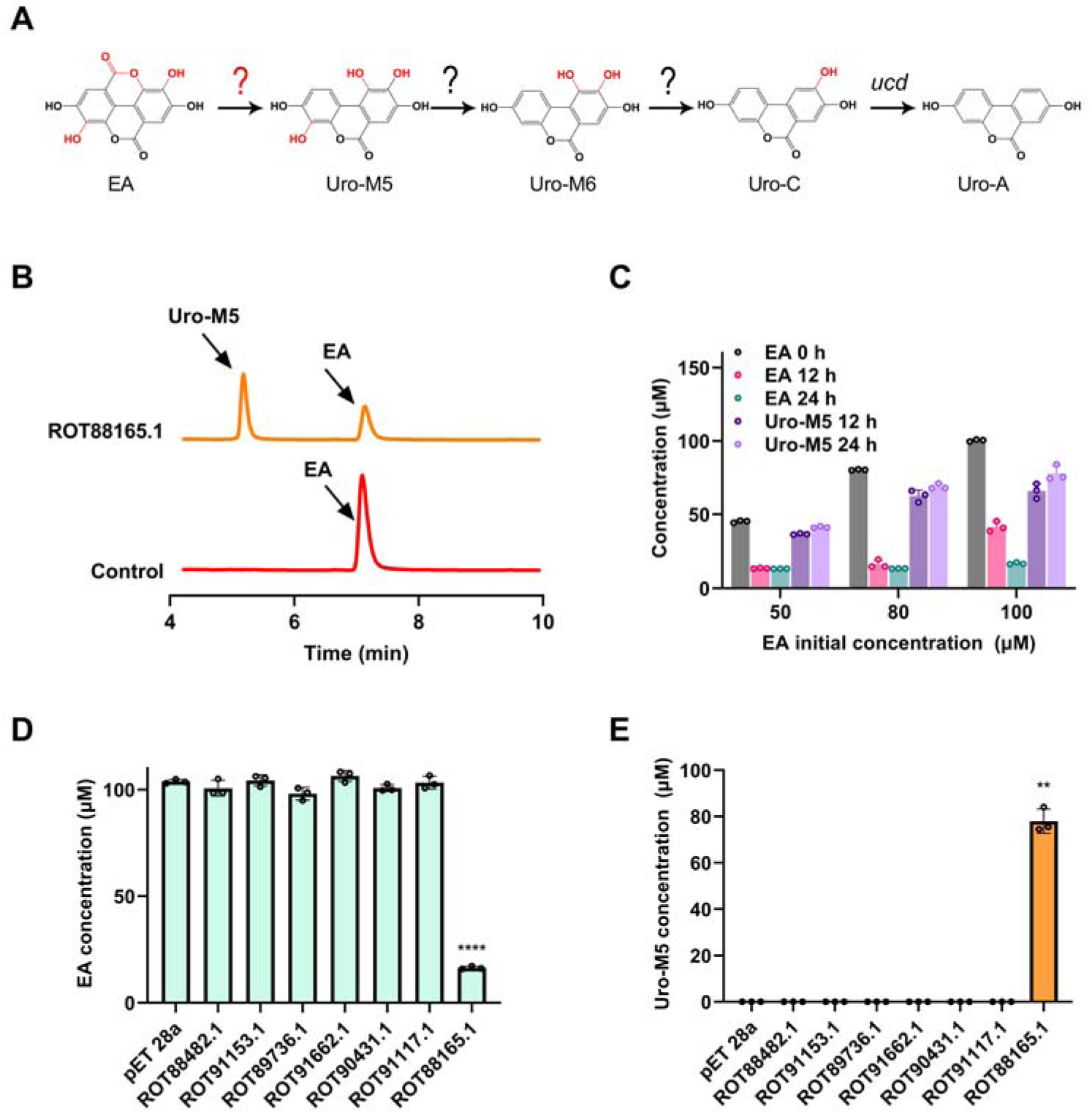
Discovery of the EAL in *G. urolithinfaciens* DSM 27213. (A) Proposed major pathway for EA metabolism by the human gut microbiota. (B) The production of Uro-M5 was detected by HPLC of *E. coli* BL21 encoding the ROT88165.1 protein. (C) Changes in concentrations of EA and Uro-M5 during the fermentation of *E. coli* BL21 encoding the ROT88165.1 protein over a 24-h period. (D) EA consumption by *E. coli* BL21 strains expressing the predicted enzymes. (E) Uro-M5 production by *E. coli* BL21 strains expressing the predicted enzymes. Data are presented as mean ± standard deviation (n = 3). Statistical significance was determined using a two-tailed Student’ s t-test (\*\**P* < 0.01, \*\*\*\**P* < 0.0001).

### Biochemical assays on the recombinant EAL

Under optimized conditions with an isopropyl β-D-1-thiogalactopyranoside (IPTG) concentration of 0.10 mM, the EAL protein was successfully expressed in *E. coli* BL21 cells through recombinant expression (Fig. 3A). A hexahistidine tag was incorporated to facilitate purification, which was subsequently performed using immobilized metal affinity chromatography with Ni-NTA resin (Fig. 3B). Ultraviolet-visible (UV) spectral analysis of the purified enzyme revealed a characteristic protein absorption profile, showing maximal absorbance at 280 nm without detectable peaks in the visible wavelength range (Fig. 3C). This finding suggests that no UV absorbing cofactors were associated with the purified EAL. Based on this, a reaction system was established without the addition of any UV absorbing cofactors, leading to a significant reduction in EA concentration and a concurrent rise in Uro-M5 production (Fig. 3D). The ability of EAL to metabolize EA into Uro-M5 (the main molecular ion fragments m/z 229.0 and 275.0) was further confirmed through UPLC-Q-TOF/MS^2^ analysis (Fig. 3E and F).

**Fig. 3.**
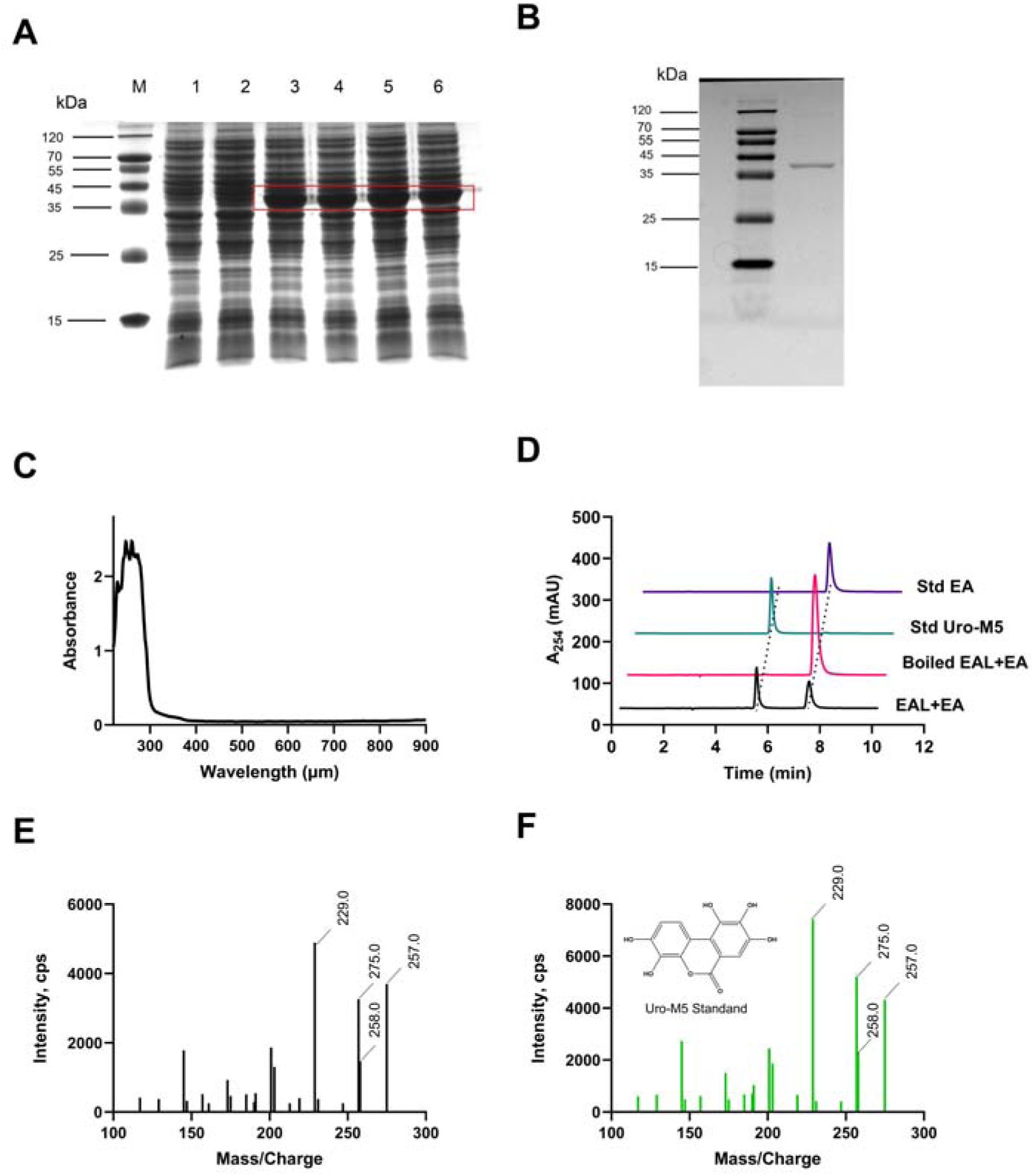
Purification and functional validation of EAL. (A) SDS-PAGE analysis of *E. coli* BL21 crushing supernatant before and after induction with different concentrations of IPTG. M, protein markers. lane 1, pET-28a(+)-IPTG (0.1 mM). lane 2, pET-28a(+)-*eal*. lane 3, pET-28a(+)-*eal*-IPTG (0.05 mM). lane 3, pET-28a(+)-*eal*-IPTG (0.10 mM). lane 5, pET-28a(+)-*eal*-IPTG (0.15 mM). lane 6, pET-28a(+)*-eal*-IPTG (0.20 mM). (B) SDS-PAGE of the purified EAL protein. (C) UV spectra of the purified EAL protein. (D) HPLC analysis of EA products catalyzed by EAL *in vitro*. Boiled EAL was used for control reactions. (E) Tandem mass spectrometry analysis of the HPLC fractions corresponding to the main product formed by incubation of EAL with EA *in vitro*. (F) Tandem mass spectrometry analysis of the of the Uro-M5 standard.

Based on the above results, we further examined how temperature and pH influence EAL activity, identifying 37°C and pH 8.5 as the optimal reaction conditions (Fig. 4A and B). Under optimal conditions (37°C, pH 8.5), we evaluated the temperature and pH stability of EAL, the impact of metal ions on its activity, its substrate specificity, and its kinetic parameters. EAL exhibited stability at temperatures up to 40°C and within a pH range of 7.0 to 9.0 (Fig. 4C and D). While Na⁺and K⁺ had no impact on EAL activity, Ca²⁺, Mg²⁺ and Ba²⁺ markedly enhanced its activity (Fig. 4E). To assess the substrate specificity of EAL, we tested 14 compounds containing linear and cyclic ester bonds, as detailed in Fig. S3. HPLC analysis revealed that recombinant EAL did not exhibit activity toward the ester bonds of these compounds, indicating a strong specificity of EAL for EA. Additionally, we determined the kinetic parameters of EAL in the conversion of EA. The reaction exhibited a typical hyperbolic curve of product formation versus substrate concentration when varying EA concentrations were used in the presence of 1 mg/mL EAL (Fig. 4F), consistent with Michaelis-Menten kinetics. Quantification of Uro-M5 by HPLC allowed us to estimate the apparent steady-state kinetic constants. Nonlinear regression analysis yielded the following parameters: *K*_m_ = 0.3305 ± 0.115 mM and *V*_max_ = 0.5975 ± 0.08 μmol·s⁻¹·mg⁻¹. The above results indicate that EA has a high specific affinity for EAL.

**Fig. 4.**
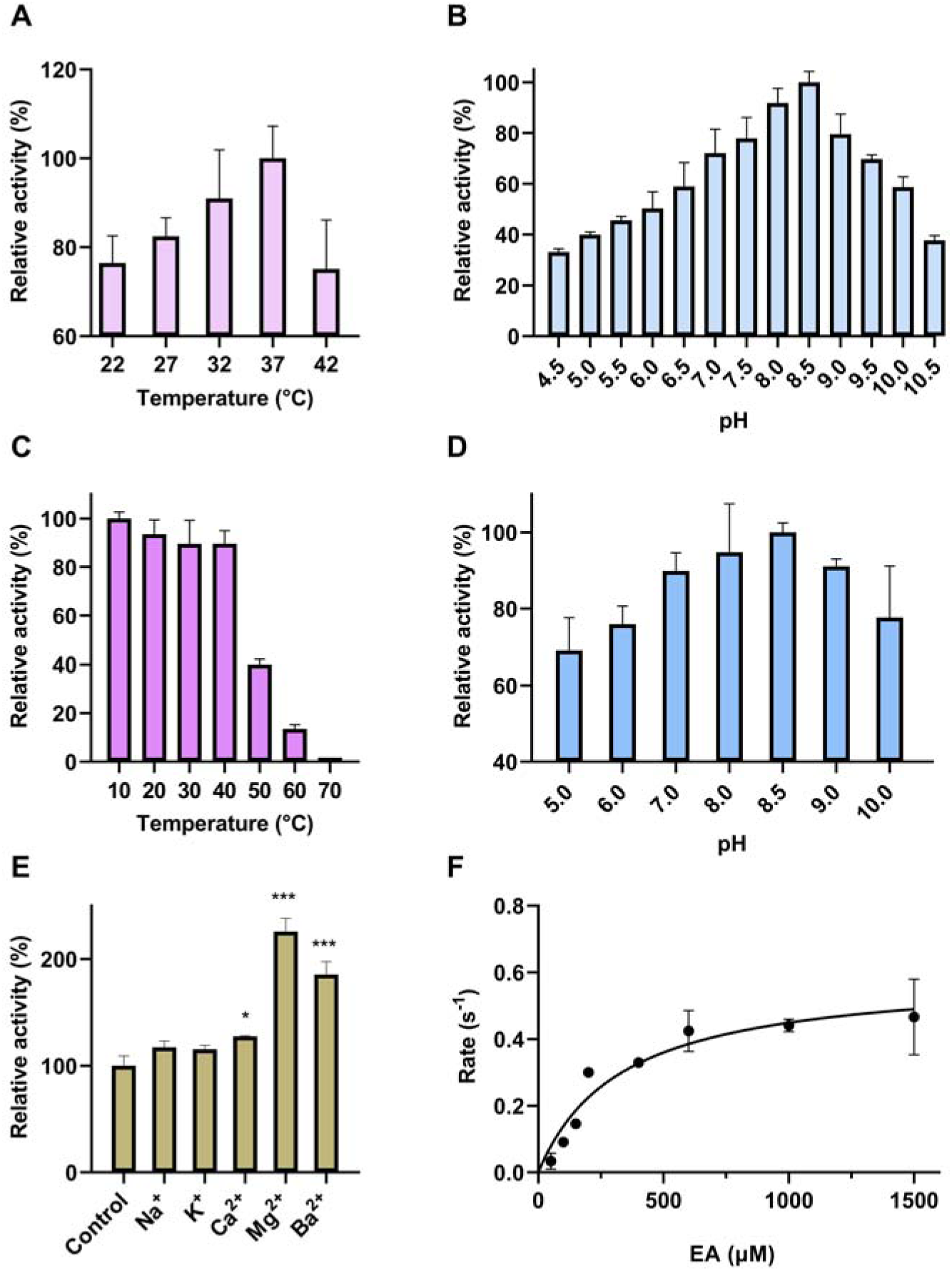
Biochemical assays of EAL. (A) Dependency of the EAL activity on temperature. (B) Dependency of the EAL activity on pH. (C) Stability of EAL against temperature. (D) Stability of EAL against pH. (E) Effects of metals ions on the activity of EAL. (F) Michaelis-Menten plot of the EA conversion reaction catalyzed by EAL. Results are presented as mean ± standard deviation (n = 3). Statistical significance was assessed using a two-tailed Student’s t-test (\**P* < 0.05, \*\*\**P* < 0.001).

### Phylogenetic analysis of EAL

We utilized the EAL protein sequence from *G. urolithinfaciens* DSM 27213 as a query to investigate the distribution of EAL-like enzymes across microbial communities. A BLASTP search identified 865 putative EAL enzymes (amino acid sequence identity ≥35%, coverage ≥80%, e-value ≤1e^−10^) from over 330 microbial species, indicating a widespread presence of these enzymes among microorganisms. To elucidate their phylogenetic relationships, we performed a maximum likelihood phylogenetic analysis of 100 putative EAL enzymes from more than 90 microbial strains (Table S5). The analysis revealed that EAL homologues are present in both aerobic and anaerobic bacteria from aquatic, terrestrial, and human digestive environments. Most EALs were found in prokaryotes, with significant proportions present in primary gut bacterial groups such as Actinomycetota and Bacillota (Fig. 5A), this distribution aligns with the observation that many EA-degrading bacteria are intestinal actinomycetes (19). Additionally, EALs were also detected in a few members of Alphaproteobacteria, Gammaproteobacteria, Chloroflexota, Archaea, Desulfobacteria, and Planctomycetota.

**Fig. 5.**
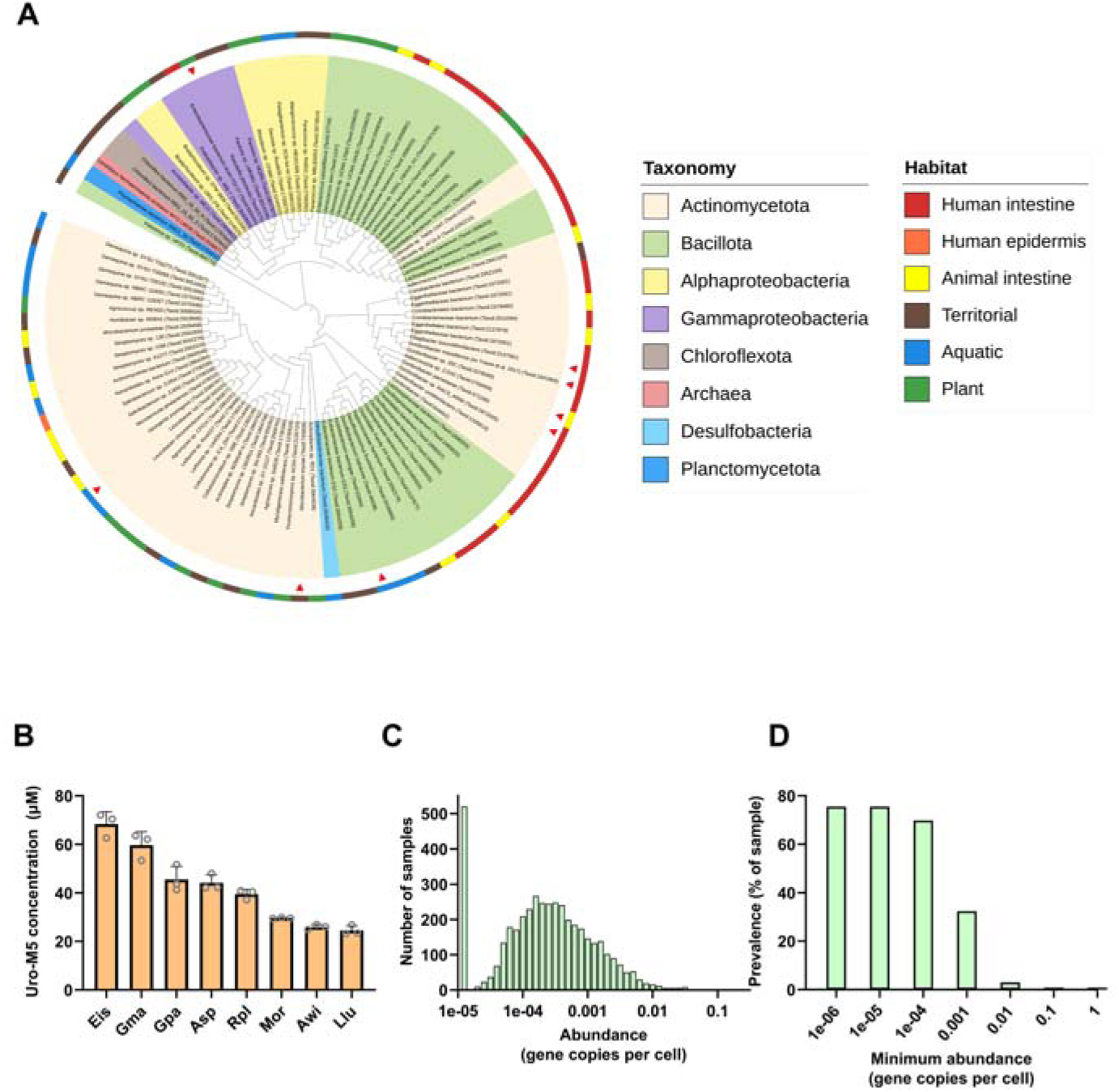
Distribution of EAL. (A) Phylogenetic analysis of EALs. EAL-like enzyme labeled with red triangle was selected to determine its activity against EA. (B) The activity of EAL homologues from different microbial hosts was analyzed using EA as substrate. (C) The abundance of EAL homologues in human gut microbiome samples. (D) The prevalence of EAL homologues in human gut metagenome under different abundance thresholds. Results are presented as mean ± standard deviation (n = 3).

To validate the phylogenetic analysis, eight putative EALs from different isolates were selected following a strategy of decreasing sequence similarity (Table S6). These candidates were then tested for their metabolic activity on EA. As anticipated, all tested enzymes exhibited EA conversion activity (Fig. 5B), supporting the reliability of the EALs identified in the phylogenetic tree.

We further investigated the distribution and abundance of EAL-like enzymes within the human gut microbiota using the MetaQuery database. The analysis revealed that putative *eal* genes are widespread in the human gut microbiota, with abundances ranging from 0.0001 (one copy per 10,000 cells) to 0.01 (one copy per 100 cells) (Fig. 5C). The average estimated copy number of the gene is approximately one per 1,000 cells. Additionally, we assessed the distribution of the *eal* gene across different abundance thresholds in the human gut metagenome, revealing its presence in 69.80% of samples, with a minimum abundance of 0.0001 (one copy per 10,000 cells) (Fig. 5D). These results indicate that EALs are widely present in the human microbial community, although further investigations are needed to confirm their degradation activity.

### Sequence similarity network (SSN) analysis of EAL

The EALs identified in this study, based on their catalytic properties, are classified as carboxylester hydrolases (CEHs, EC 3.1.1.-). CEHs are enzymes that hydrolyze both linear and cyclic ester bonds to yield carboxyl (-COOH) and hydroxyl (-OH) groups at the termini, and they play a role in the transformation of phenolic acids (20). To systematically evaluate the relationship between EALs and known cyclic esterases, we constructed an SSN utilizing the 100 EAL sequences (Table S5) identified in this study and approximately 1,506 sequences of known cyclic esterases from seven families (Table S1) in the UniProtKB protein database. The SSN analysis revealed that EALs constitute a highly diverse group of lactonases that remain largely uncharacterized (Fig. 6A).

**Fig. 6.**
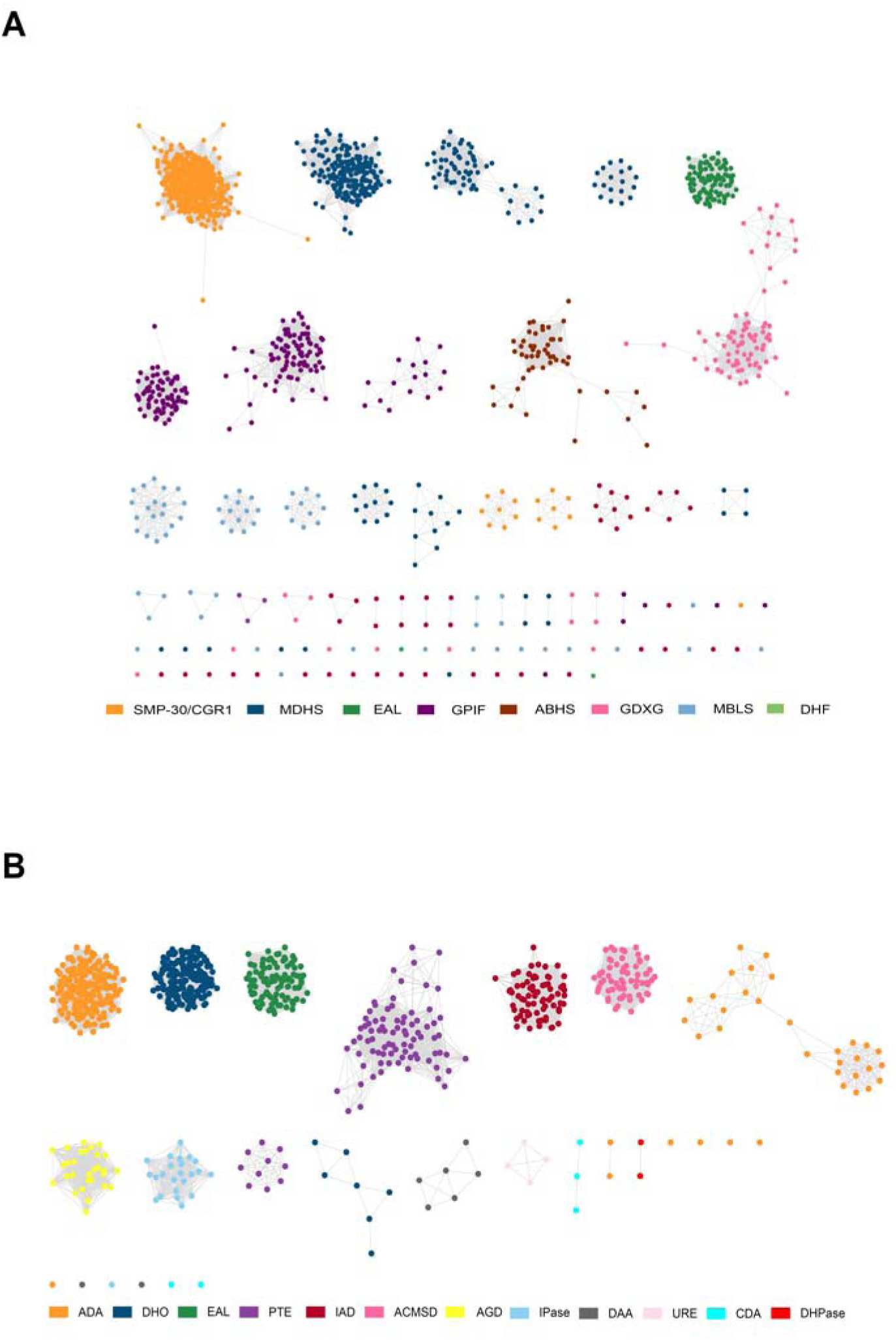
SSN analysis of EAL. (A) Construction of SSN for lactonases. Each node is colored according to cluster type. (B) Construction of amidohydrolase family SSN. Each node is colored according to cluster type. Enzymes are color-coded according to their functional classification, forming discrete clusters that potentially group proteins with analogous catalytic properties. Within the phylogenetic tree, 100 EAL-like proteins are highlighted in green to represent their evolutionary relationships.

Additionally, we noted that the target protein ROT88165.1 is annotated as an amidohydrolase with both carboxyl-lyase and hydrolase activities. The amidohydrolase superfamily consists of over five subfamilies, and, similar to α/β-hydrolases, amidohydrolases are capable of catalyzing various hydrolysis reactions, typically involving ester and amide bonds (21). To further investigate the relationship between EALs and known amidohydrolases, we constructed an SSN that included 11 well-characterized members of the amidohydrolase family (Fig. 6B). The SSN analysis indicated that EALs do not fall within the established categories of amidohydrolases but represent a novel subset within this enzymatic family.

### Structural prediction and catalytic mechanism of EAL

The above studies have validated the function of EAL. Consequently, we downloaded the predicted crystal structure using AlphaFold2 from the AlphaFold Protein Structure Database (Fig. 7A), aiming to elucidate the detailed catalytic mechanism of EAL. The accuracy of the predicted structures was assessed using the Template Modeling (TM) score, calculated by ResQ (https://seq2fun.dcmb.med.umich.edu//ResQ/), with scores ranging from 0 to 1.0, where higher scores correspond to greater accuracy (22). The TM score of 0.79 predicted by AlphaFold2 for the EAL structure indicates that the structural model has good reliability. We proceeded with the structural analysis based on the AlphaFold2 prediction. As depicted in Fig. 7A, EAL adopts a central (α/β)7 barrel structure, with seven β-strands forming the core and seven α-helices encircling the exterior. This arrangement creates a distinct, thermally stable pocket near the active site, characterized by a propeller-like shape due to the β-strand configuration. The overall structure aligns with the typical features observed in the amidohydrolase superfamily (23). Substrate specificity is primarily governed by structural elements on the catalytic face of the (α/β)_7/8_ barrel, including loops, insertions, and conformational variation, which aligns with the observed substrate specificity of EAL (Fig. S3). Additionally, size-exclusion chromatography-multi-angle light scattering (SEC-MALS) analysis confirmed the functional unit of EAL as a homotetramer (Fig. 7B and Fig. S4).

**Fig. 7.**
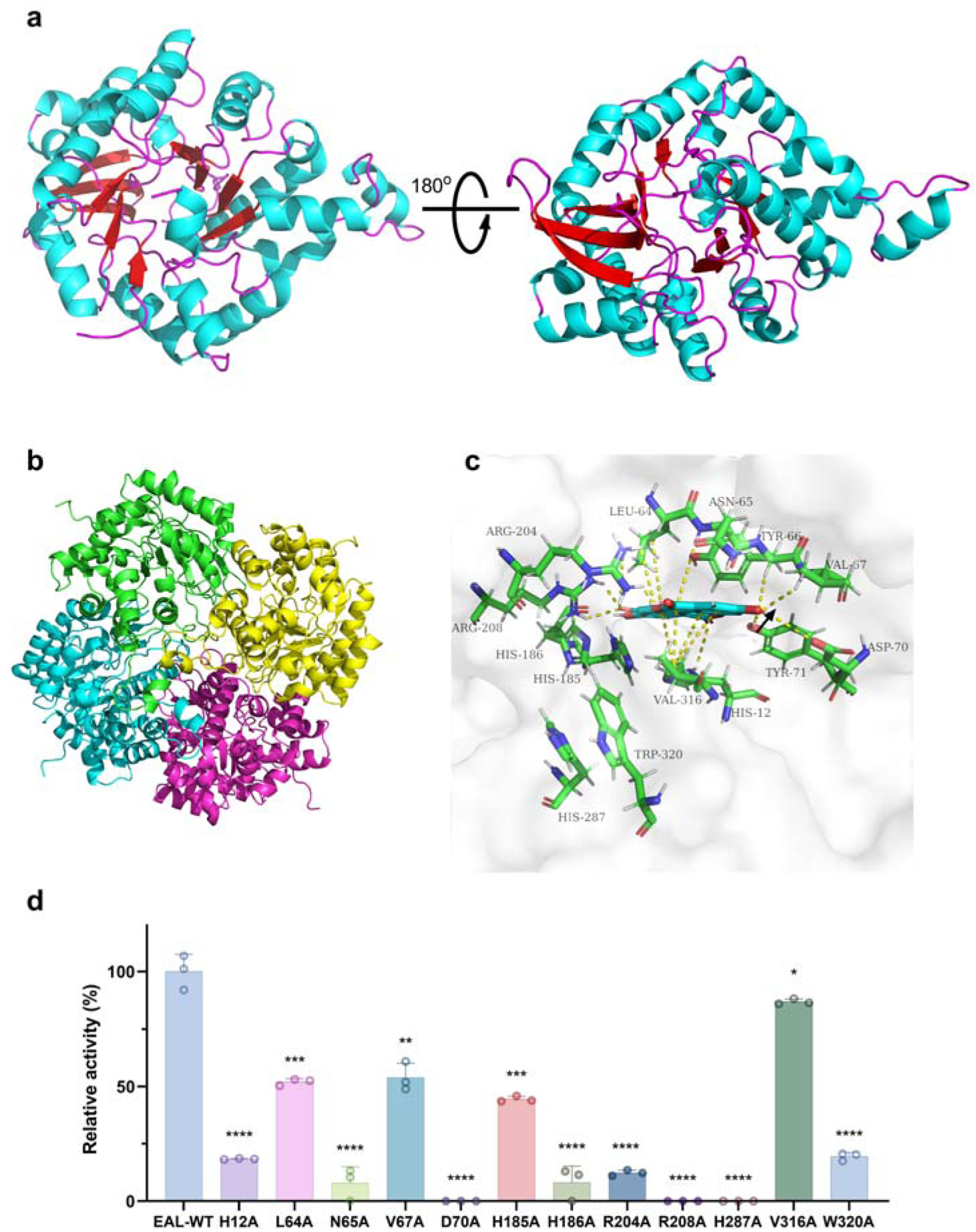
Structural analysis of EAL. (A) The predicted EAL monomer structure by Alphfold2. (B) Homotetramer structure of EAL predicted by ZDOCK. (C) Zoom-in views of the EAL and EA-binding site. (D) Assay of enzyme activity of wild-type or EAL mutants *in vitro*. Results are presented as mean ± standard deviation (n = 3). Statistical significance was assessed using a two-tailed Student’s t-test (\**P* < 0.05. \*\**P* < 0.01 \*\*\**P* < 0.001. \*\*\*\**P*< 0.0001).

To further investigate the catalytic mechanism of EAL on EA, we employed molecular docking techniques to simulate the binding of EA to EAL (Fig. 7C). As anticipated, EA was positioned within the active pocket of EAL. The molecular dynamics simulation results, including the root mean square fluctuation (RMSF), root mean square deviation (RMSD), radius of gyration (Rg) and solvent accessible surface area (SASA), demonstrated the stability of the protein-ligand complex. The low RMSF and stable RMSD values (within 0.1–0.3 nm) indicated minimal structural fluctuations and a well-maintained conformation. Additionally, the consistent Rg and SASA values suggested a compact structure and stable solvent accessibility, further supporting the reliability of the docking results (24) (Fig. S5).The key residues involved in substrate binding include H12, Y66, V67, D70, and Y71, which form hydrogen bonds with EA, while R204 and R208 engage in both electrostatic and hydrogen bonding interactions. Additionally, L64 and V316 exhibit hydrophobic interactions, and N65 shows mixed interactions with EA. To explore the reaction mechanism of EAL, a series of single amino acid substitution mutants were constructed: H12A, L64A, N65A, H185A, H186A, R204A, R208A, H287A, V316A, and V320A, targeting residues within the active pocket that are strictly conserved (Fig. S6). Each mutant was expressed and purified in *E. coli* BL21 according to the protocol outlined in the Materials and Methods. Catalytic activity of the mutants was markedly lower than that of the wild type, with the R208A and H287A mutants completely abolishing the hydrolytic activity of EAL (Fig. 7D). In summary, this study deepens our understanding of the enzymatic function of EAL and the dynamics of substrate interactions.

## Discussion

The metabolism of physiologically active compounds such as phenolic compounds, flavonoids, alkaloids, and steroids by microorganisms is a complex process that remains partially understood. Recent studies have identified several microbial metabolic enzymes responsible for the transformation of these compounds (25-31). Despite this, our understanding of the molecular components driving gut microbiota functions remains incomplete. This study identified EALs as a previously unrecognized class of lactonases/amidohydrolases that played a key role in the conversion of EA to Uro-M5. Our findings shed light on the early stages of EA conversion at both the gene and protein levels, advancing our understanding of Uro-A production in the human body. This discovery lays a foundation for future research into the relationship between Uros and human health.

Biochemical assays of EAL confirmed that the enzyme exhibited optimal activity at 37°C, with over 70% activity at a pH of 7.0-7.5, conditions typical of the human gut (32). This suggested that EAL could specifically bind to EA within the intestinal microenvironment and facilitate its conversion to Uro-M5. Notably, EAL displayed a strict substrate specificity, not metabolizing any of the linear and cyclic ester compounds we tested, despite the structural similarity of the daphnetin compound to EA. This may be attributed to the fact that EA is a polyphenolic dilactone containing two six-membered rings (δ-lactone). The high stability of the δ-lactone rings likely contributes to the strict substrate specificity of EAL (33). Kinetic studies demonstrated that EAL follows Michaelis-Menten kinetics, with parameters indicating a strong affinity for EA. The high conversion efficiency of EA to Uro-M5 in the presence of recombinant EAL expressed in *E. coli* BL21 further underscores its potential utility in biotechnological applications aimed at utilizing EA in metabolic processes (Fig. 2C).

Lactones, commonly found in nature, are integral to various biological processes (34). Enzymes involved in lactone metabolism have been partially identified, but detailed investigations of these enzymes are still lacking (35, 36). Lactonases are known to influence physiological processes by metabolizing and modifying the biological activity of both endogenous and exogenous lactones (37). EA is one of the most abundant lactones found in nature (38). Although we identified EAL through sequence analysis of known lactonases, subsequent SSN analysis indicated that EAL and its homologs differ from previously known lactonases. The protein (ROT88165.1) encoded by EAL is annotated in the UniProt database as an amidohydrolase with dual hydrolase and carboxyl-lyase activities. Our results indicated that EAL represents a novel addition to the amidohydrolase superfamily. This superfamily includes a variety of enzymes that catalyze the hydrolysis of substrates containing amide or ester groups at carbon and phosphorus centers, including urease (URE) (39), dihydroorotase (DHO) (40), and isoaspartyl dipeptidase (IAD) (41), which cleave C-N bonds, as well as phosphotriesterase (PTE), which hydrolyze phosphoester bonds in organophosphorus triesters (42). While EALs share characteristics with known amidohydrolases, they constitute a unique subset within this family, reflecting their distinct evolutionary pathways.

The amidohydrolase superfamily is defined by a conserved (β/α)7/8-barrel structural motif, one of the most common protein folds, representing around 10% of all structurally characterized proteins (21). SEC-MALS demonstrated that EAL from gut bacteria functions as a homotetramer, in contrast to microbial lactonases from *Fusarium oxysporum*, *Agrobacterium tumefaciens*, and *Acinetobacter calcoaceticus*, which predominantly exist as dimers (32). Moreover, a defining feature of these enzymes is their metal dependency (23). Our study confirmed that EAL conformed to these patterns, with divalent metal ions Ca²⁺, Mg²⁺, and Ba²⁺ significantly enhancing its activity (Fig. 4E). Additionally, it was noted that other divalent metal ions such as Zn²⁺, Ni²⁺, and Fe²⁺ could also augment the activity of related amidohydrolases (21). Within the amidohydrolase superfamily, the fifth β-strand typically provides His ligands for metal coordination, while the His residue on the sixth β-strand can serve either as a metal ligand or a non-ligand, playing a critical role in the active site architecture (21). Docking studies further revealed that the conserved catalytic His residues in EAL (His12, His185, His186, and His287) interact with EA (Fig. 7D). Site-directed mutagenesis confirmed that these conserved amino acids are essential for the enzymatic activity of EAL, opening potential avenues for enzyme engineering aimed at enhancing substrate specificity or activity.

Our findings suggested that a variety of microorganisms, particularly those inhabiting the gut, utilized EAL to metabolize EA into Uro-M5. This widespread presence implied that EAL homologues had evolved across diverse environmental contexts, underscoring the ecological significance of EA degradation in microbial communities. Analysis of the MetaQuery database of the human microbiome confirmed that EAL-like genes were not only abundant but also widely distributed within the human gut microbiota. The presence of the eal gene in this microbial community was likely attributed to horizontal gene transfer, yet the underlying factors driving its specific distribution patterns remained unclear.

In conclusion, our study identified EAL as a key enzyme responsible for metabolizing EA into Uro-M5 and highlighted its ecological relevance within the human gut microbiome. Further research was required to elucidate the functional roles of EALs in various microbial communities and their potential implications for human health, particularly regarding dietary polyphenol metabolism. The insights gained here provided a valuable foundation for further research into the biotechnological applications of EAL and its homologues, as well as a deeper understanding of gut microbiota interactions.

## Conclusions

In this study, we identified and characterized an EAL from the gut bacterium *G. urolithinfaciens* DSM 27213, a key enzyme responsible for the initial metabolic breakdown of EA, uncovering critical insights into the enzymatic mechanisms driving the production of Uro-A in the human gut. This discovery not only enhances our understanding of the microbial pathways involved in polyphenol metabolism but also highlights the intricate interplay between gut microbiota and dietary compounds, with broader implications for human health and nutrition. By clarifying the enzymatic basis of Uro-A production, this research provides a foundation for developing targeted strategies to modulate gut microbiota activity, potentially enhancing the bioavailability and efficacy of dietary polyphenols.

## Methods

### Bacterial strains and chemicals

The strain *G. urolithinfaciens* DSM 27213, acquired from the German Collection of Microorganisms and Cell Cultures (DSMZ, Braunschweig, Germany), was cultivated in modified Gifu anaerobic medium. For plasmid construction and recombinant protein expression, *E. coli* DH5α and BL21 (DE3) strains, provided by Sangong (Shanghai, China), were used. Molecular biology reagents, ApexHF HS DNA Polymerase FS Master Mix and the Steady Pure Agarose Gel DNA Purification Kit (Accurate Biotechnology (Hunan) Co., Ltd., China) were employed. Restriction enzymes and DNA ligase, were obtained from Takara Biotechnology Co., Ltd. (Dalian, China), and oligonucleotide primers were custom-synthesized by Sangong (Shanghai, China). All anaerobic procedures were performed using a COY Vinyl Anaerobic Chamber (Coy Laboratory Products Inc., Grass Lake, MI, USA) under controlled atmospheric conditions (80% N_2_, 10% H_2_, and 10% CO_2_) at a constant temperature of 37°C.

### Molecular cloning, heterologous expression, and protein purification of candidate EAL

The potential *eal* gene from *G. urolithinfaciens* DSM 27213 was PCR-amplified using the primers listed in Table S7. The purified PCR fragment was digested with *Nhe*I and *Hin*dIII, subsequently cloned and inserted into the linearized pET-28a(+) vector, and transformed into *E. coli* DH5α competent cells to obtain the recombinant plasmids. Following sequencing verification, the constructed plasmid was introduced into *E. coli* BL21 for expression screening. The *E. coli* BL21 transformants were subsequently grown in LB supplemented with 50 µg/mL kanamycin at 37°C. Upon reaching an OD600 of approximately 0.8, enzyme expression was induced by adding IPTG to a final concentration of 0.1 mM, along with EA at 0.1 mM. Following 24 h of incubation, the culture was centrifuged at 5000×g for 10 min at 4°C, and the supernatant was subjected to HPLC analysis to quantify Uro-M5. Recombinant strains confirmed to be successful were subsequently purified.

The recombinant *E. coli* BL21 strain was plated on LB agar supplemented with 50 μg/mL kanamycin for single colony isolation. The selected colonies were grown in 10 mL of antibiotic-containing LB medium at 37°C for 16 h. The overnight culture was then transferred to 1 L of fresh LB medium with the same antibiotic concentration and incubated at 37°C until the mid-log phase (OD600 ≈ 0.8). Protein expression was induced by adding IPTG to a final concentration of 0.1 mM, followed by an 8 h incubation at 37°C. After induction, the cells were harvest via low-temperature centrifugation (5000×g, 30 min, 4°C). The bacterial pellet was then resuspended in binding buffer containing 0.5 M NaCl and 20 mM Tris·HCl (pH 8.0). Cellular disruption was achieved using ultrasonic homogenization (Ningbo Xinzhi Biotechnology Co., Ltd., China). The lysate was subjected to identical centrifugation conditions for clarification, and the resulting supernatant was membrane-filtered through a 0.45 μm pore size filter before purification. For protein purification, the clarified supernatant was subjected to immobilized metal affinity chromatography using Ni2+-Sepharose™ 6 Fast Flow resin (GE Healthcare, USA). Column equilibration was achieved by washing with 10–15 volumes of binding buffer until the UV absorbance reached equilibrium. The target proteins were eluted via a linear gradient of imidazole (0–0.5 M) in binding buffer, with fraction collection based on UV absorbance peaks. The eluted protein was concentrated and desalted via a 30 kDa molecular weight cutoff membrane, followed by the addition of a cryoprotectant (5% glycerol) before storage at -80°C. Protein quality and quantity were assessed by SDS-PAGE analysis and the Bradford protein assay, respectively.

### Quantification of EA and Uro-M5

Quantitative analysis of EA and Uro-M5 was performed on an Agilent 1200 HPLC-DAD system (Agilent Technologies, Japan) equipped with a ZORBAX Eclipse XDB-C18 reversed-phase column (250 mm × 4.6 mm *i.d.*, 5 µm particle size). A binary solvent system comprising 0.1% phosphoric acid (solvent A) and acetonitrile (solvent B) was employed with the following gradient program: isocratic at 20% A for 10 min, followed by a step increase to 30% A at 10.1 min, which was maintained until 15 min. Chromatographic detection was performed at 254 nm with a constant flow rate of 1.0 mL/min and an injection volume of 10 µL. Quantification was achieved by establishing standard calibration curves through peak area integration.

### UPLC-Q-TOF/MS^2^ measurements

Chromatographic separation was performed on a SCIEX ExionLC™ system equipped with a Kinetex XB-C18 column (250 mm × 4.6 mm *i.d.*, 5 µm particle size). The binary mobile phase consisted of 0.1% aqueous formic acid (solvent A) and acetonitrile (solvent B), and was delivered at a constant flow rate of 0.5 mL/min, and the column temperature was maintained at 40°C. The gradient program was optimized as follows: 50% B (0-2 min), a linear increase to 70% B (2-8 min), isocratic elution at 70% B (8-11 min), a stepwise increase to 90% B (11-13 min), maintenance for 15 min, and then return to the initial conditions (15-15.01 min). Mass spectrometric detection was carried out via an AB SCIEX X500R QTOF instrument with electrospray ionization in negative ion mode for qualitative analysis. Multiple reaction monitoring high-resolution (MRMHR) sampling mode was used for quantification, with the following optimal mass spectrometry parameters: the scanning mode was set to MRM, the spray voltage was -4500 V, the ion source temperature was 550°C, the curtain gas pressure was set to 35 psi, the collision gas pressure was set to 9 psi, and the nebulizer gas (GS1) and auxiliary gas (GS2) pressures was set to to 55 psi. Data acquisition and analysis were performed via SCIEX OS 1.7.2 software. To maintain the precision and dependability of the quantification process, each batch of analysis included quality control standards.

### Enzymatic properties of EAL

The pH optimization for EAL activity was conducted by incubating 0.1 mM EA with 10 μL of purified recombinant enzyme (protein concentration: 1.0 mg/mL) in 20 mM buffer systems across different pH ranges: phosphate-citrate buffer (pH 4.5-6.0), potassium phosphate buffer (pH 6.5-8.0), and Gly-NaOH buffer (pH 8.5-10.5). For pH dependency estimations, the residual enzyme activity of recombinant EAL treated overnight at 4°C in different pH buffers was assessed under optimal conditions.

To establish the optimal assay parameters for EAL, enzymatic reactions were conducted at 22, 27, 32, 37, and 42°C for 10 min. For thermal dependency estimation, the enzyme mixture was pretreated at temperatures ranging from 10 to 70°C (in 10°C increments) for 1 h, followed by activity measurement under the established optimal assay conditions.

The effects of divalent and monovalent cations (including Na⁺, K⁺, Ca²⁺, Mg²⁺, and Ba²⁺ in the form of chloride salts) on EAL activity were evaluated at a fixed concentration of 5 mM. Additionally, substrate specificity was assessed by screening a panel of structurally diverse compounds, including chlorogenic acid, rosmarinic acid, chicoric acid, ethyl caffeate, scoparone, psoralen, scopoletin, dihydrocoumarin, bergapten, imperatorin, daphnetin, umbelliferone, atractylenolide I, and atractylenolide III. All the compounds were tested at a concentration of 0.1 mM.

The enzymatic reactions were terminated by heating at 80°C for 5 min, followed by quantification of the generated products via HPLC. All the experiments were conducted in an anaerobic environment and in triplicate.

### Enzyme activity assay

Enzyme activity was assessed in a reaction system with a total volume of 1 mL, comprising 10 μL of enzyme mixture (protein concentration: 1.0 mg/mL) incubated with 900 μL of alkaline buffer (20 mM Gly-NaOH, pH 8.5). The reaction was started by adding EA to a final concentration of 100 μM and incubated anaerobically at 37°C for 10 min. After termination by heating at 80°C for 5 min, the samples were mixed with an equal volume of methanol and centrifuged at 5000×g for 5 min. The supernatant was subsequently subjected to HPLC analysis to determine the concentrations of EA and Uro-M5. Each experiment was conducted in triplicate.

### Kinetic parameter determination

Enzyme kinetics were determined by monitoring the initial rate of the reaction with varying concentrations of EA (50-1500 μM) via HPLC. Enzymatic reactions were performed in a total volume of 1 mL, consisting of 900 μL of alkaline buffer (20 mM Gly-NaOH, pH 8.5), 10 μL of enzyme solution (1.0 mg/mL protein), and 90 μL of substrate solution, and incubated anaerobically at 37°C. Each reaction was repeated in triplicate. The kinetic constants (*K*_m_ and *V*_max_) were calculated by performing nonlinear regression analysis of the experimental data using the Michaelis-Menten model, as implemented in GraphPad Prism software.

### Molecular docking and molecular dynamic simulations

The docking of proteins and ligands was performed using the CDOCKER module in Discovery Studio 2021. The protein structures used in this study were obtained from the AlphaFold Protein Structure Database (43). The 3D molecular structures of all the compounds were retrieved from the PubChem database. Prior to docking analysis, the protein was preprocessed by removing water molecules and adding hydrogen atoms, and small molecules that had undergone structural optimization were selected as ligands. The interactions between the protein and small molecules post-docking were visualized using the View Interactions tools module in Discovery Studio, and the 3D structure of the protein-ligand complex was visualized using PyMOL.

The EAL-EA complex was subjected to a 50-ns molecular dynamics simulation employing GROMACS 2018, utilizing the CHARMM27 force field within an SPC216 water model in a cubic periodic boundary condition. The v-rescale algorithm was employed to maintain a constant temperature, while the Berendsen algorithm was used for pressure control. Prior to the simulation, energy minimization and NVT (canonical ensemble) and NPT (isothermal-isobaric ensemble) equilibrations were performed. After trajectory file analysis, the RMSF, RMSD, Rg and SASA curves were generated as a function of time for further analysis.

### Phylogenetic investigation of EAL homologs

The EAL protein sequence (GenBank: ROT88165.1) was used as a query to perform a BLASTp search against the NCBI non-redundant protein database. The top 100 homologous sequences with the highest similarity scores were retrieved and aligned using Clustal W (version 2.1) under default parameters. The multiple sequence alignment was subsequently analyzed in MEGA X (version 10.0.13) to construct a phylogenetic tree (44). Phylogenetic reconstruction was performed using the maximum-likelihood method based on the Le-Gascuel (LG) substitution model, which was determined by the maximum-likelihood criterion. The reliability of the phylogenetic tree was evaluated through bootstrap analysis with 1000 replicates to provide statistical support for branch confidence. The consensus tree was further displayed and annotated using the web-based program iTOL.

### Analysis of EAL homolog abundance in the human gut microbiota

To assess the prevalence of EAL homologs within the human gut microbiome, computational analysis was conducted using the MetaQuery web-based platform (https://metaquery.docpollard.org/) (45). The alignment parameters were configured as follows: minimum percent identity >40%, maximum E-value <1e^−05^, and percent coverage of query with minimum query/target alignment coverage >70%.

### Construction and functional annotation of an SSN

The SSN was generated based on protein sequences of amidohydrolases and lactonases utilizing the Enzyme Function Initiative online tool (available at http://efi.igb.illinois.edu/efi-est/) (46). The alignment parameters were set with an alignment score of 30, a maximum sequence length of 400, and a minimum sequence length of 200. The resulting network was visualized and annotated using Cytoscape 3.10.2 (47). The amidohydrolases analyzed included adenosine deaminase (ADA), DHO, PTE, IAD, α-amino-β-carboxymuconate-ε-serialdehyde decarboxylase (ACMSD), N-acetylglucosamine 6-phosphate deacetylase (AGD), imidazolone propionase (IPase), D-aminoacetylase (DAA), URE, cytosine deaminase (CDA) and dihydropyrimidinase (DHPase).

The lactonase family encompasses several groups, including the SMP-30/CGR1 family (SMP-30/CGR1), the metal-dependent hydrolases superfamily (MDHS), the metal β-lactamase superfamily (MBLS), the glucose/galactosamine-6-phosphate isomerase family (GPIF), the α/β-hydrolases superfamily (ABHS), the dienolactone hydrolase family (DHF), and the ‘GDXG’ lipid enzyme family (GDXG).

### Site-directed mutagenesis

To investigate the functional roles of key residues within the active site of EAL, site-directed mutagenesis was performed to substitute individual residues with alanine using the QuickMutation™ Site-Directed Gene Mutagenesis Kit (Beyotime Biotechnology Co., Ltd., Shanghai, China). The recombinant pET-28a(+)-*eal* plasmid served as the template for these mutations, utilizing the primers listed in Table S7. The resulting EAL variants were transformed into *E. coli* DH5α cells. The expression, purification, and enzyme assay conditions for these variants were conducted under conditions identical to those used for the wild-type enzyme.

### Analysis of SEC-MALS

SEC-MALS was performed using a TSKgel G3000SWXL column (7.8 mm × 300 mm i.d., 5 µm particle size) The column was equilibrated at 25°C with a buffer containing 200 mM NaLJHPOLJ and 1% isopropanol (pH 7.0). A 100 μL EAL solution (∼2 mg/mL) was injected, and the buffer flow rate was set to 0.5 mL/min. Data were recorded and analyzed using ASTRA 7.3.1.9 software (Wyatt Technologies, USA).

### Statistical analysis

For all the data, the *P*-values were calculated using a two-tailed Student’s t-test, and a *P*-value of ≤ 0.05 was considered statistically significant.

## Data availability

Data will be made available on request.

## Author contributions

ZX conceived and designed and directed the study, isolated the gene, constructed the plasmid, identified the enzyme, purified the enzyme, and constructed a sequence similarity network, analyzed the catalytic mechanism and wrote the manuscript. JY constructed the evolutionary tree and verified the enzyme substrate specificity. FX contributed to this study by participating in the construction of plasmids, generating site-directed mutations, and characterizing the enzymatic properties. FC and QM participated in protein purification and analysis of enzymatic properties. YG participated in the characterization of enzymatic properties. ZR supervised the study, and edited the manuscript. All authors read and approved the final manuscript.

