## Supplemental Material for "Discovery and characterization of a lactonase in gut microbiota that initiates the metabolism of ellagic acid"

### **Supplementary Tables**

Table S1. Known lactone hydrolases and their docking interactions with EA.

| Famliy | Lactase name | EC number |
| --- | --- | --- |
| MDHS | **Phospho-furanose lactonase** | EC 3.1.1.104  EC 3.1.1.25 |
|  | **L-rhamnono-gamma-lactonase** | EC:3.1.1.65 |
|  | **D-arabinonolactonase** | EC 3.1.1.30 |
|  | **4-sulfomuconolactone hydrolase** | EC 3.1.1.92 |
|  | **2-pyrone-4,6-dicarbaxylate hydrolase** | EC 3.1.1.57 |
| SMP-30/CGR1 | **L-arabinolactonase** | EC:3.1.1.15 |
|  | **Gluconolactonase** | EC:3.1.1.17 |
|  | Pentonolactonase XacC | EC 3.1.1.15  EC 3.1.1.68 |
|  | **Xylono-1,4-lactonase** | EC 3.1.1.68 |
|  | **6-deoxy-6-sulfogluconolactonase** | EC:3.1.1.99 |
| MBLS | **N-acyl homoserine lactonase** | EC:3.1.1.81 |
|  | **4-pyridoxolactonase** | EC 3.1.1.27 |
| GPIF | **6-phosphogluconolactonase** | EC 3.1.1.31 |
| ABHS | Dihydrocoumarin hydrolase | EC:3.1.1.35 |
|  | 3-oxoadipate enol-lactonase | EC 3.1.1.24 |
| DHF | **2-oxo-3-(5-oxofuran-2-ylidene) propanoate lactonase** | EC:3.1.1.91 |
| GDXG | **Monoterpene epsilon-lactone hydrolase** | EC:3.1.1.83 |

MDHS, metallo-dependent hydrolases superfamily. SMP-30/CGR1, SMP-30/CGR1 family. MBLS, metallo-beta-lactamase superfamily. GPIF, glucosamine/galactosamine-6-phosphate isomerase family. ABHS, AB hydrolase superfamily. DHF, dienelactone hydrolase family. GDXG, 'GDXG' lipolytic enzyme family.

Table S2. Docking parameter of currently known lactonases.

| Lactase name | BINDING ENERGY (kcal/mol) | -CDOCKER ENERGY (kcal/mol) | -CDOCKER INTERACTION ENERGY (kcal/mol) |
| --- | --- | --- | --- |
| **Phospho-furanose lactonase** | -442.595 | 65.93 | 76.838 |
| **6-phosphogluconolactonase** | -423.854 | 68.112 | 78.895 |
| **6-deoxy-6-sulfogluconolactonase** | -199.77 | 44.714 | 59.127 |
| **L-rhamnono-gamma-lactonase** | -183.35 | 52.097 | 52.883 |
| **Gluconolactonase** | -145.203 | 40.745 | 41.705 |
| **4-sulfomuconolactone hydrolase** | -89.416 | 43.025 | 43.566 |
| Pentonolactonase XacC | -76.886 | 33.026 | 43.074 |
| **2-oxo-3-(5-oxofuran-2-ylidene) propanoate lactonase** | -65.975 | 32.701 | 34.554 |
| **L-arabinolactonase** | -55.748 | 8.536 | 22.217 |
| **2-pyrone-4,6-dicarbaxylate hydrolase** | -32.636 | 23.147 | 32.074 |
| **N-acyl homoserine lactonase** | -26.842 | 26.691 | 31.42 |
| Dihydrocoumarin hydrolase | -25.787 | 25.662 | 27.288 |
| **Xylono-1,4-lactonase** | -24.385 | 27.146 | 38.332 |
| **D-arabinonolactonase** | -0.017 | 28.35 | 27.829 |

Table S3. Potential EAL identified through DELTA-BLAST in *G. urolithinfaciens* DSM 27213.

| Name | Protein IDs | Gene IDs | Uniprot IDs |
| --- | --- | --- | --- |
| lactonase family protein | ROT91662.1 | DMP12_03205 | A0A423UNA5 |
| gluconolactonase | ROT91236.1 | DMP12_04160 | A0A423UML6 |
| amidohydrolase | ROT91117.1 | DMP12_03695 | A0A423UMA7 |
| amidohydrolase | ROT88165.1 | DMP12_13210 | A0A423UH90 |
| TatD family deoxyribonuclease | ROT89090.1 | DMP12_10490 | A0A423UIU5 |
| hypothetical protein | ROT89966.1 | DMP12_07590 | A0A423UKD5 |
| peptidase | ROT88897.1 | DMP12_10600 | A0A423UIN5 |
| 4Fe-4S dicluster domain-containing protein | ROT91153.1 | DMP12_03900 | A0A423UMC9 |
| putative heme d1 biosynthesis radical SAM protein NirJ2 | ROT88482.1 | DMP12_12230 | A0A423UHU4 |
| glucosamine-6-phosphate deaminase | ROT89736.1 | DMP12_08305 | A0A423UJY6 |
| acetyl esterase | ROT90431.1 | DMP12_05280 | A0A423UL04 |
| alpha/beta hydrolase | ROT88342.1 | DMP12_12330 | A0A423UHJ9 |
| DUF2974 domain-containing protein | ROT92136.1 | DMP12_01210 | A0A423UP39- |
| CocE/NonD family hydrolase | ROT88153.1 | DMP12_13150 | A0A423UH94 |
| alpha/beta hydrolase | ROT89108.1 | DMP12_10455 | A0A423UIZ6 |
| MBL fold metallo-hydrolase | ROT89054.1 | DMP12_10245 | A0A423UIP9- |
| MBL fold metallo-hydrolase | ROT88497.1 | DMP12_12295 | A0A423UHU2 |
| MBL fold metallo-hydrolase | ROT90594.1 | DMP12_05395 | A0A423ULJ1 |
| hypothetical protein | ROT88359.1 | DMP12_12425 | A0A423UHQ3 |
| ribonuclease J | ROT88679.1 | DMP12_11495 | A0A423UI81 |
| MBL fold metallo-hydrolase | ROT89368.1 | DMP12_09155 | A0A423UJ92 |
| FprA family A-type flavoprotein | ROT89707.1 | DMP12_08160 | A0A423UJW1 |
| alpha/beta hydrolase | ROT88025.1 | DMP12_13710 | A0A423UGY8 |
| alpha/beta hydrolase | ROT90012.1 | DMP12_07865 | A0A423UKG5 |
| alpha/beta hydrolase | ROT89973.1 | DMP12_07640 | A0A423UKF0 |
| alpha/beta hydrolase | ROT89108.1 | DMP12_10455 | A0A423UIZ6 |
| alpha/beta hydrolase | ROT88342.1 | DMP12_12330 | A0A423UHJ9 |

Table S4. Docking parameter of potential EAL.

| Name | Protein IDs | BINDING ENERGY (kcal/mol) | -CDOCKER ENERGY (kcal/mol) | -CDOCKER INTERACTION ENERGY (kcal/mol) |
| --- | --- | --- | --- | --- |
| acetyl esterase | ROT90431.1 | -341.707 | 39.381 | 58.72 |
| glucosamine-6-phosphate deaminase | ROT89736.1 | -323.713 | 38.911 | 49.911 |
| putative heme d1 biosynthesis radical SAM protein NirJ2 | ROT88482.1 | -321.195 | 44.021 | 56.187 |
| 4Fe-4S dicluster domain-containing protein | ROT91153.1 | -267.588 | 40.67 | 52.145 |
| amidohydrolase | ROT88165.1 | -251.406 | 2.516 | 33.853 |
| amidohydrolase | ROT91117.1 | -230.852 | 29.087 | 39.756 |
| TatD family deoxyribonuclease | ROT89090.1 | -207.519 | 8.476 | 34.98 |
| DUF2974 domain-containing protein | ROT92136.1 | -204.117 | 37.785 | 37.242 |
| alpha/beta hydrolase | ROT88025.1 | -190.924 | 37.524 | 48.608 |
| alpha/beta hydrolase | ROT88342.1 | -187.712 | 39.824 | 51.546 |
| alpha/beta hydrolase | ROT88342.1 | -187.712 | 39.824 | 51.546 |
| MBL fold metallo-hydrolase | ROT88497.1 | -183.283 | 28.573 | 38.739 |
| MBL fold metallo-hydrolase | ROT89054.1 | -183.257 | 32.23 | 43.487 |
| ribonuclease J | ROT88679.1 | -151.919 | 17.307 | 28.087 |
| lactonase family protein | ROT91662.1 | -133.122 | 32.254 | 32.108 |
| hypothetical protein | ROT89966.1 | -119.634 | 17.144 | 33.016 |
| hypothetical protein | ROT88359.1 | -111.848 | 22.258 | 32.88 |
| alpha/beta hydrolase | ROT90012.1 | -109.614 | 0.493 | 20.381 |
| MBL fold metallo-hydrolase | ROT90594.1 | -95.241 | 14.847 | 29.052 |
| gluconolactonase | ROT91236.1 | -85.747 | 26.568 | 29.353 |
| MBL fold metallo-hydrolase | ROT89368.1 | -48.107 | 3.641 | 18.148 |
| peptidase | ROT88897.1 | -42.555 | 1.601 | 27.801 |

Table S5. 100 EALs utilized for constructing phylogenetic evolutionary trees.

| Description | Per. ident | Accession |
| --- | --- | --- |
| *Acetobacterium wieringae* | 99.72 | WP_301156404.1 |
| *Adlercreutzia* sp. ZJ242 | 99.72 | WP_087190618.1 |
| *Agromyces* sp. CF514 | 98.01 | WP_015539932.1 |
| *Bradyrhizobium* sp. STM 3809 | 95.74 | WP_165170380.1 |
| *Candidatus Nezhaarchaeota* archaeon WYZ-LMO8 | 82.67 | WP_114602621.1 |
| *Cellulosimicrobium* sp. I38E | 81.82 | WP_221688628.1 |
| *Chloroflexi bacterium* RBG_16_50_9 | 80.68 | MDD5806909.1 |
| *Clostridiales* bacterium | 78.98 | WP_158048450.1 |
| *Collinsella* sp. TM09-10AT | 78.98 | MDO4290986.1 |
| *Coriobacteriia* bacterium | 78.12 | PWL78437.1 |
| *Demequina* sp. SYSU T00192 | 77.01 | WP_251212307.1 |
| *Desulfosporosinus fructosivorans* | 74.71 | HAM14591.1 |
| *Eggerthellales* bacterium | 73.30 | MDR0515335.1 |
| *Ellagibacter isourolithinifaciens* | 72.78 | MBQ2682165.1 |
| *Enterobacteriaceae* bacterium strain FGI 57 | 71.84 | MDR3307946.1 |
| *Eubacteriaceae* bacterium ES2 | 63.40 | WP_148638568.1 |
| *Gordonibacter massiliensis* (ex Traore et al. 2017) | 63.11 | WP_135552428.1 |
| *Gordonibacter pamelaeae* | 62.54 | MCL1896220.1 |
| *Gordonibacter* sp. 28C | 62.25 | WP_015943688.1 |
| *Gordonibacter* sp. RACS_AR68 | 62.07 | MDR3288029.1 |
| *Gordonibacter urolithinfaciens* | 61.38 | WKY45144.1 |
| *Lachnospiraceae* bacterium | 60.86 | MCD7902642.1 |
| *Leifsonia* sp. Root227 | 60.81 | WKY46794.1 |
| *Leucobacter luti* | 60.63 | MCD7934552.1 |
| *Microbacterium oryzae* | 59.37 | MCF8095320.1 |
| *Myceligenerans salitolerans* | 59.20 | WP_290106647.1 |
| *Nocardioides* sp. Kera G14 | 59.20 | WP_136316968.1 |
| *Oenococcus* sp. UCMA 17063 | 58.05 | WP_143624211.1 |
| *Oscillospiraceae* bacterium | 57.93 | MCD8004553.1 |
| *Pantoea* sp. MQR6 | 57.88 | WP_207276054.1 |
| *Paracoccus* sp. PAR01 | 57.76 | WP_227781660.1 |
| *Pelosinus* sp. UFO1 | 57.47 | MDQ4114282.1 |
| *Peptococcaceae* bacterium | 57.35 | MDR3304535.1 |
| *Pokkaliibacter* sp. MBI-7 | 57.31 | WP_143417095.1 |
| *Promicromonospora* sp. AC04 | 57.18 | WP_307533990.1 |
| *Raoultella planticola* | 56.73 | WP_263509603.1 |
| *Ruminococcus* sp. D55t1_190419_H1 | 56.73 | WP_136571031.1 |
| *Salinibacterium* sp. ZJ454 | 56.61 | WP_064316720.1 |
| *Streptomyces* sp. A1277 | 56.61 | WP_055889237.1 |
| *Acinetobacte*r sp. ME22 | 56.61 | WP_306207098.1 |
| *Actinomycetota* bacterium | 56.45 | WP_104474789.1 |
| *Actinoplanes* sp. RD1 | 56.32 | WP_092963952.1 |
| *Actinotalea* sp. M2MS4P-6 | 56.32 | WP_130453156.1 |
| *Adlercreutzia murintestinalis* | 56.32 | WP_056731432.1 |
| *Agrococcus* sp. REN33 | 56.32 | WP_166937549.1 |
| *Agromyces* sp. Soil535 | 56.03 | WP_167044643.1 |
| *Bradyrhizobium* sp. ORS 278 | 56.03 | WP_055901943.1 |
| *Candidatus Stoquefichus* sp. SB1 | 56.03 | WP_202345494.1 |
| *Cellulomonas* sp. IC4_254 | 55.75 | WP_108491557.1 |
| *Chloroflexi* bacterium RBG_16_50_9 | 55.75 | WP_166879740.1 |
| *Clostridiales* bacterium TF09-2AC | 55.59 | WP_202232796.1 |
| Clostridiales Family XIII bacterium | 55.46 | WP_301134466.1 |
| *Collinsella* sp. AF16-8 | 55.17 | WP_084099545.1 |
| *Coriobacteriaceae* bacterium | 55.17 | WP_301159963.1 |
| *Coriobacteriales* bacterium | 55.17 | WP_073922423.1 |
| *Demequina* sp. NBRC 110051 | 55.04 | WP_084159777.1 |
| *Demequina* sp. NBRC 110057 | 54.89 | WP_301153457.1 |
| *Demequina* sp. SYSU T00068 | 54.60 | WP_306231595.1 |
| *Demequina* sp. SYSU T0a273 | 54.60 | WP_243065150.1 |
| *Desulfitobacterium hafniense* | 54.31 | WP_084079157.1 |
| *Desulfobacteraceae* bacterium | 51.72 | WP_129388010.1 |
| *Devosia* sp. Root635 | 48.99 | MCD8128916.1 |
| *Eggerthellaceae* bacterium | 48.41 | MBE6016240.1 |
| *Eggerthellaceae* bacterium | 45.40 | WP_008966382.1 |
| *Eggerthellaceae* bacterium | 44.99 | WP_243080034.1 |
| *Enterocloster clostridioformis* | 44.41 | WP_032699882.1 |
| *Enterococcus casseliflavus* | 44.29 | KXT72536.1 |
| *Erysipelotrichaceae* bacterium | 44.25 | WP_007757695.1 |
| *Eubacteriaceae* bacterium ES3 | 44.16 | WP_011925428.1 |
| *Georgenia yuyongxinii* | 44.13 | WP_269295136.1 |
| *Humibacter* sp. RRB41 | 43.97 | WP_279789669.1 |
| *Lachnospiraceae* bacterium | 43.55 | AGB77295.1 |
| *Lachnospiraceae* bacterium | 43.55 | ODT83677.1 |
| *Lachnospiraceae* bacterium | 43.52 | MDE6202259.1 |
| *Lachnospiraceae* bacterium | 43.27 | WP_216671116.1 |
| *Lachnospiraceae* bacterium | 42.94 | WP_012962264.1 |
| *Lachnospiraceae* bacterium C1.1 | 42.86 | WP_225612247.1 |
| *Leifsonia* sp. Leaf264 | 42.69 | WP_217933828.1 |
| *Leucobacter chromiireducens* | 42.53 | WP_195476964.1 |
| *Mangrovicoccus* sp. HB161399 | 42.53 | WP_311162659.1 |
| *Microbacterium protaetiae* | 42.41 | WP_056231654.1 |
| *Microterricola pindariensis* | 41.79 | TDA36372.1 |
| *Nocardioides* sp. GY 10127 | 41.67 | WP_237815141.1 |
| *Oenococcus oeni* | 41.59 | KPK74331.1 |
| *Oenococcus* sp. UCMA 16435 | 41.09 | MCR5271755.1 |
| *Oscillospiraceae* bacterium | 40.92 | WP_050637635.1 |
| *Oscillospiraceae* bacterium | 40.80 | MDO4192243.1 |
| *Oscillospiraceae* bacterium | 40.80 | MCD8336113.1 |
| *Pantoea* sp. LMR881 | 40.52 | MCD8074801.1 |
| *Paracoccus* sp. MBLB3053 | 40.46 | MCD8053262.1 |
| *Pelagibacterium* sp. SCN 64-44 | 40.35 | MDN6968166.1 |
| *Phycisphaerae* bacterium SM23_30 | 40.35 | WP_002821215.1 |
| *Rhizobium* sp. CF080 | 40.35 | AZZ61697.1 |
| *Salinibacterium* sp. ZJ450 | 40.06 | OGO23004.1 |
| *Streptococcus gallolyticus* | 39.77 | OGO22994.1 |
| *Streptococcus gallolyticus* | 39.48 | WP_117647437.1 |
| *Streptomyces* sp. 130 | 39.48 | WP_117778910.1 |
| *Streptomyces* sp. CB03911 | 39.19 | RJW48757.1 |
| *Streptomyces* sp. SN-593 | 39.02 | MDN4743822.1 |
| *Streptomyces* sp. V3I8 | 38.04 | WP_270289997.1 |

Table S6. Eight EAL homologues from various microorganisms were used to assess the reliability of the phylogenetic evolutionary tree.

| Abbr. | Organism | Per. ident | Protein |
| --- | --- | --- | --- |
| gpa | *Gordonibacter pamelaeae* | 98.01 | Wp_015539932.1 |
| asp | *Adlercreutzia* sp. ZJ242 | 95.74 | Wp_165170380.1 |
| gma | *Gordonibacter massiliensis* | 81.82 | Wp_221688628.1 |
| eis | *Ellagibacter isourolithinifaciens* | 78.98 | Wp_158048450.1 |
| awi | *Acetobacterium wieringae* | 63.40 | Wp_148638568.1 |
| mor | *Microbacterium oryzae* | 59.20 | Wp_290106647.1 |
| llu | *Leucobacter luti* | 56.32 | Wp_130453156.1 |
| rpl | *Raoultella planticola* | 44.41 | Wp_032699882.1 |

Table S7. Primers used in this study.

| Primer name | Sequence (5′-3′) |
| --- | --- |
| DMP12_12230-F (NdeI) | CGCCATATGatgctcgtgtcgtggatgac |
| DMP12_12230-R (HindIII) | ACTAAGCTTctagacggcagccgctac |
| DMP12_08305-F (NdeI) | CGCCATATGatggaattcatcatcgccgag |
| DMP12_08305-R (HindIII) | CCCAAGCTTttacgccaaatcgcggcag |
| DMP12_03205-F (NdeI) | CGCCATATGatgaaaggagtctgcgcc |
| DMP12_03205-R (HindIII) | CCCAAGCTTctacccgtcccagatgacg |
| DMP12_03695-F (NdeI) | CGCCATATGatgaccgccatcgacg |
| DMP12_03695-R (HindIII) | CCCAAGCTTctaaaccctcacgcccgtg |
| DMP12_03900-F (NdeI) | CGCCATATGatgaggcccgatttggaatc |
| DMP12_03900-R (HindIII) | CCCAAGCTTctactcgaagtgctcggct |
| DMP12_05280-F (NdeI) | CGCCATATGatgaacaagatcgacgtgct |
| DMP12_05280-R (HindIII) | ACTAAGCTTctgcgcctaggatggc |
| DMP12_13210-F (NdeI) | CGCCATATGatggcagacaacaaggtcat |
| DMP12_13210-R (HindIII) | CCCAAGCTTctacaggttgaacagcttcgc |
| H12A-F | GTCGGTGGGCAGGAAGGCCATGTTGATGTCGATGAC |
| H12A-R | GTCATCGACATCAACATGGCCTTCCTGCCCACCGAC |
| L64A-F | CGCCCTCCACGTAGTTCGCGATGTCCTTGCCGTCC |
| L64A-R | GGACGGCAAGGACATCGCGAACTACGTGGAGGGCG |
| N65A-F | GTCGCCCTCCACGTAGGCCAGGATGTCCTTGCC |
| N65A-R | GGCAAGGACATCCTGGCCTACGTGGAGGGCGAC |
| V67A-F | GGACATCCTGAACTACGCTGAGGGCGACTACAC |
| V67A-R | GTGTAGTCGCCCTCAGCGTAGTTCAGGATGTCC |
| D70A-F | ACTACGTGGAGGGCGCCTACACGCTGGAGACG |
| D70A-R | CGTCTCCAGCGTGTAGGCGCCCTCCACGTAGT |
| H185A-F | CTGTCCGGGCGTATGGGCGATGGCGCAGGGCA |
| H185A-R | TGCCCTGCGCCATCGCCCATACGCCCGGACAG |
| H186A-F | CCTGCGCCATCCACGCTACGCCCGGACAGAAC |
| H186A-R | GTTCTGTCCGGGCGTAGCGTGGATGGCGCAGG |
| R204A-F | GATACGGCCGAGCTCGGCTCGCAGGGGCGTGTA |
| R204A-R | TACACGCCCCTGCGAGCCGAGCTCGGCCGTATC |
| R208A-F | GGCCTGCACGTGGATAGCGCCGAGCTCGCGTC |
| R208A-R | GACGCGAGCTCGGCGCTATCCACGTGCAGGCC |
| H287A-F | GCTCCAGCTCATGGGGGCCGTCATGTCGAAGTAGAT |
| H287A-R | ATCTACTTCGACATGACGGCCCCCATGAGCTGGAGC |
| V316A-F | CCATCCACTCGTAGAACGCGGGGAACGAAGAGCC |
| V316A-R | GGCTCTTCGTTCCCCGCGTTCTACGAGTGGATGG |
| W320A-F | CACCGAGCGGGCCATCGCCTCGTAGAACACGGG |
| W320A-R | CCCGTGTTCTACGAGGCGATGGCCCGCTCGGTG |

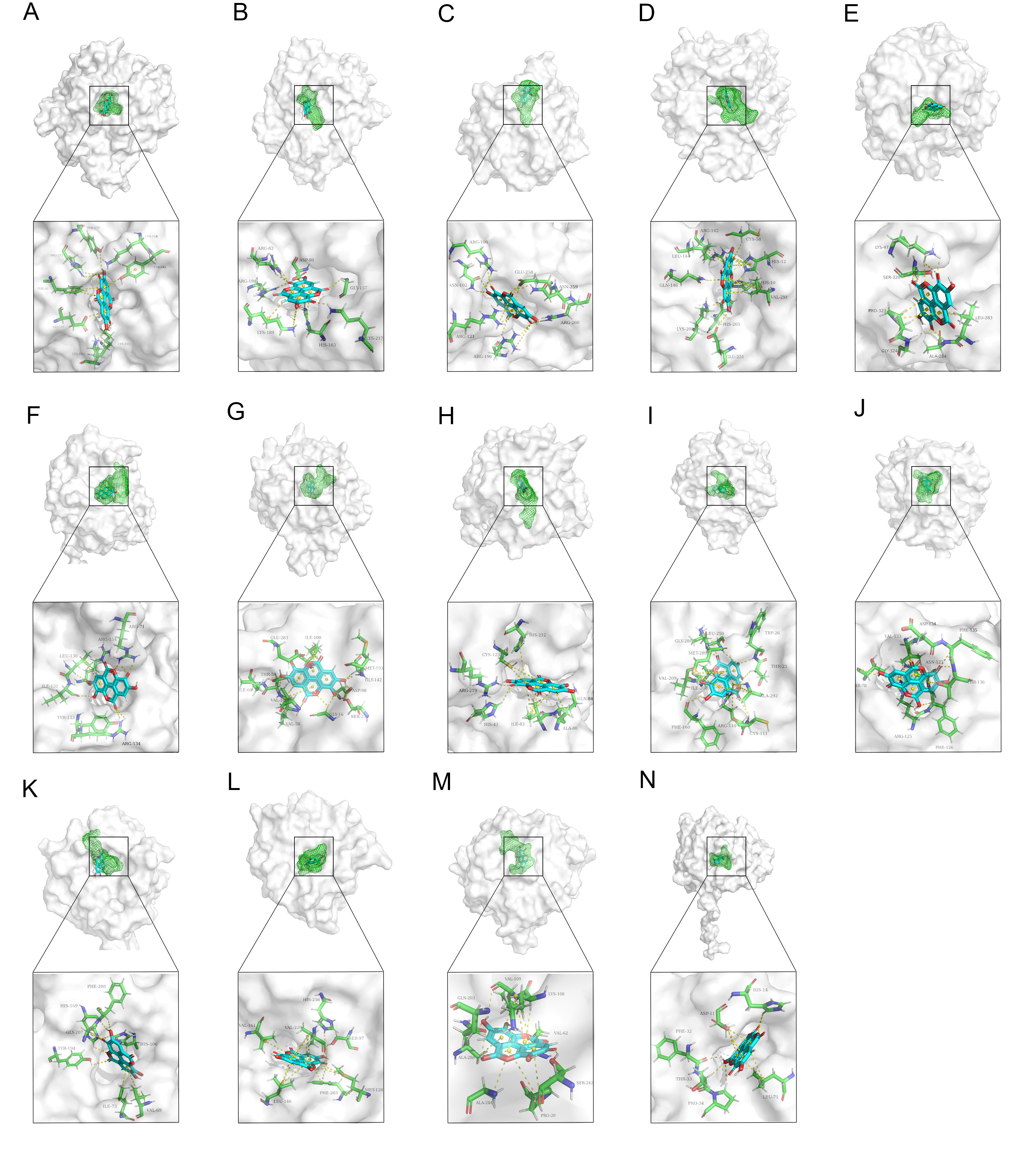

Fig. S1. Molecular docking of EA with 14 known lactonases. (A) Phospho-furanose lactonase. (B) 6-phosphogluconolactonase. (C) 6-deoxy-6-sulfogluconolactonase. (D) L-rhamnono-gamma-lactonase. (E) Gluconolactonase. (F) 4-sulfomuconolactone hydrolase. (G) Pentonolactonase XacC. (H) 2-oxo-3-(5-oxofuran-2-ylidene) propanoate lactonase. (I) L-arabinolactonase. (J) 2-pyrone-4,6-dicarboxylate hydrolase. (K) N-acyl homoserine lactonase. (L) Dihydrocoumarin hydrolase. (M) Xylono-1,4-lactonase. (N) D-arabinonolactonase.

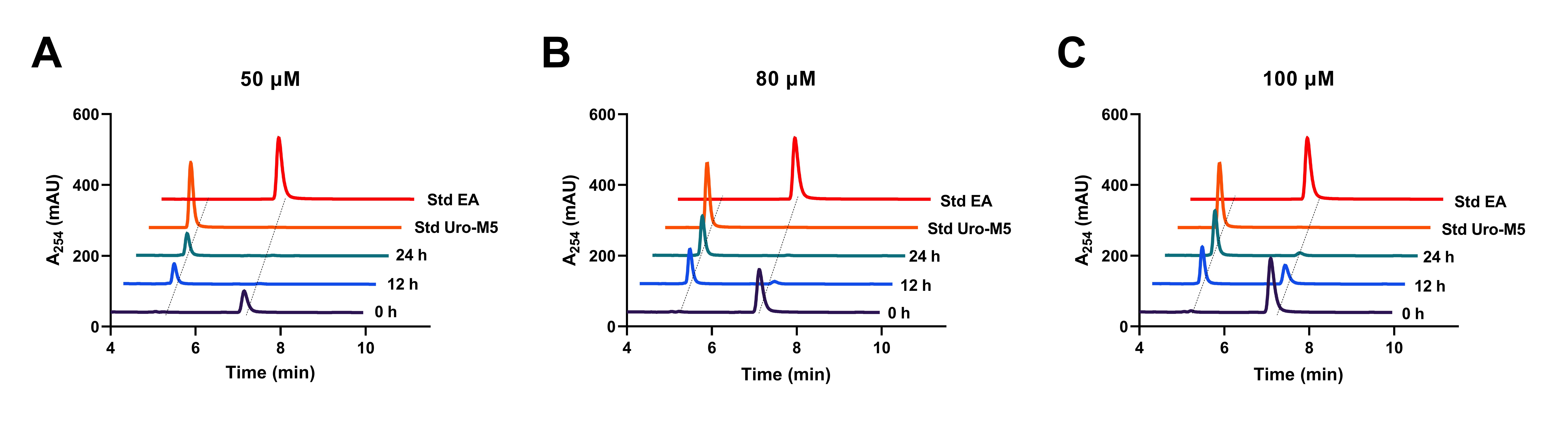

Fig. S2. The metabolic capability of *E. coli* BL21 expressing the ROT88165.1 protein convertor was assessed at varying concentrations of EA. (A) Metabolic capacity of *E. coli* BL21 containing the target protein at an EA concentration of 50 μM. (B) Metabolic capacity of *E. coli* BL21 containing the target protein at an EA concentration of 80 μM. (C) Metabolic capacity of *E. coli* BL21 containing the target protein at an EA concentration of 100 μM.

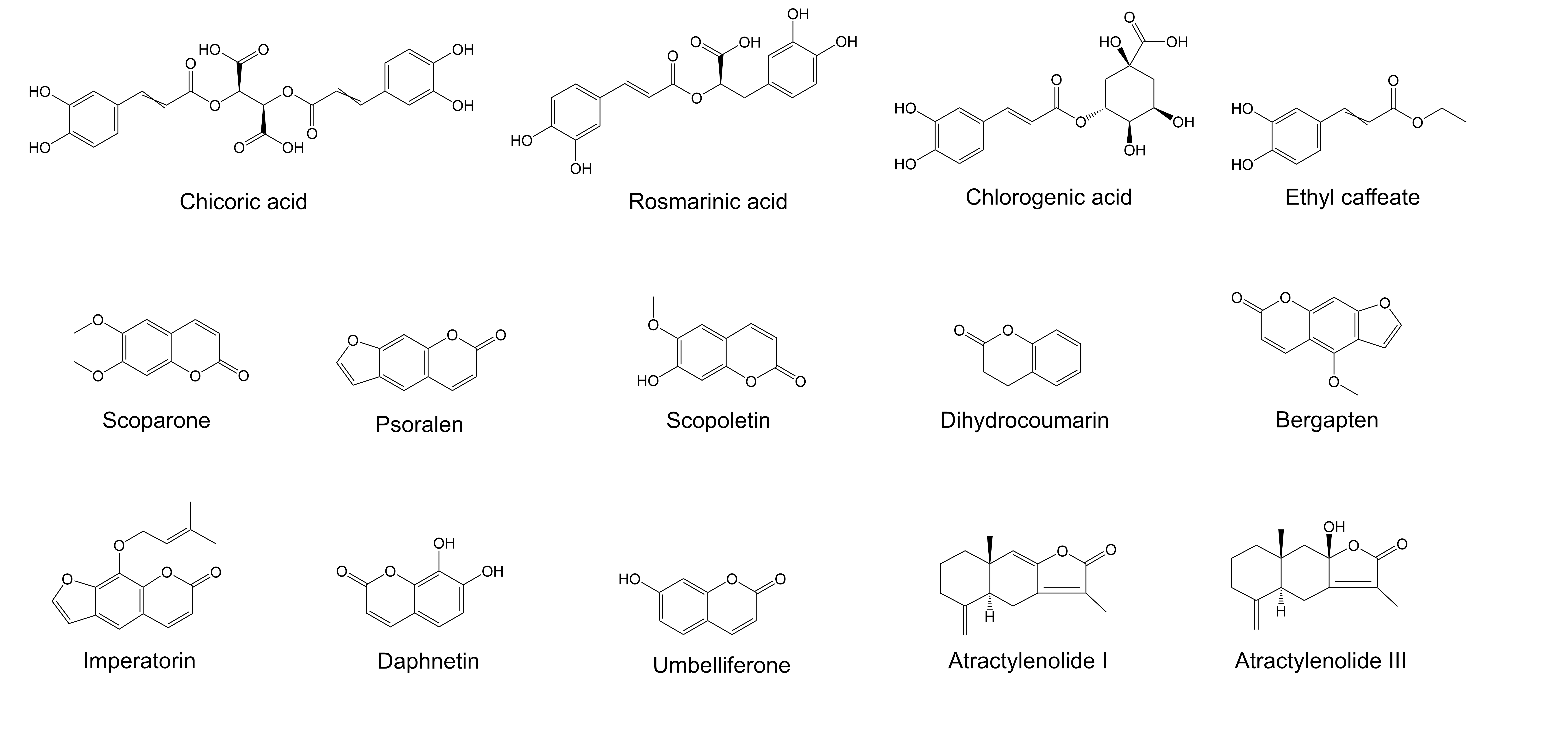

Fig. S3. Compounds containing linear and cyclic ester bonds used to validate substrate specificity of EAL.

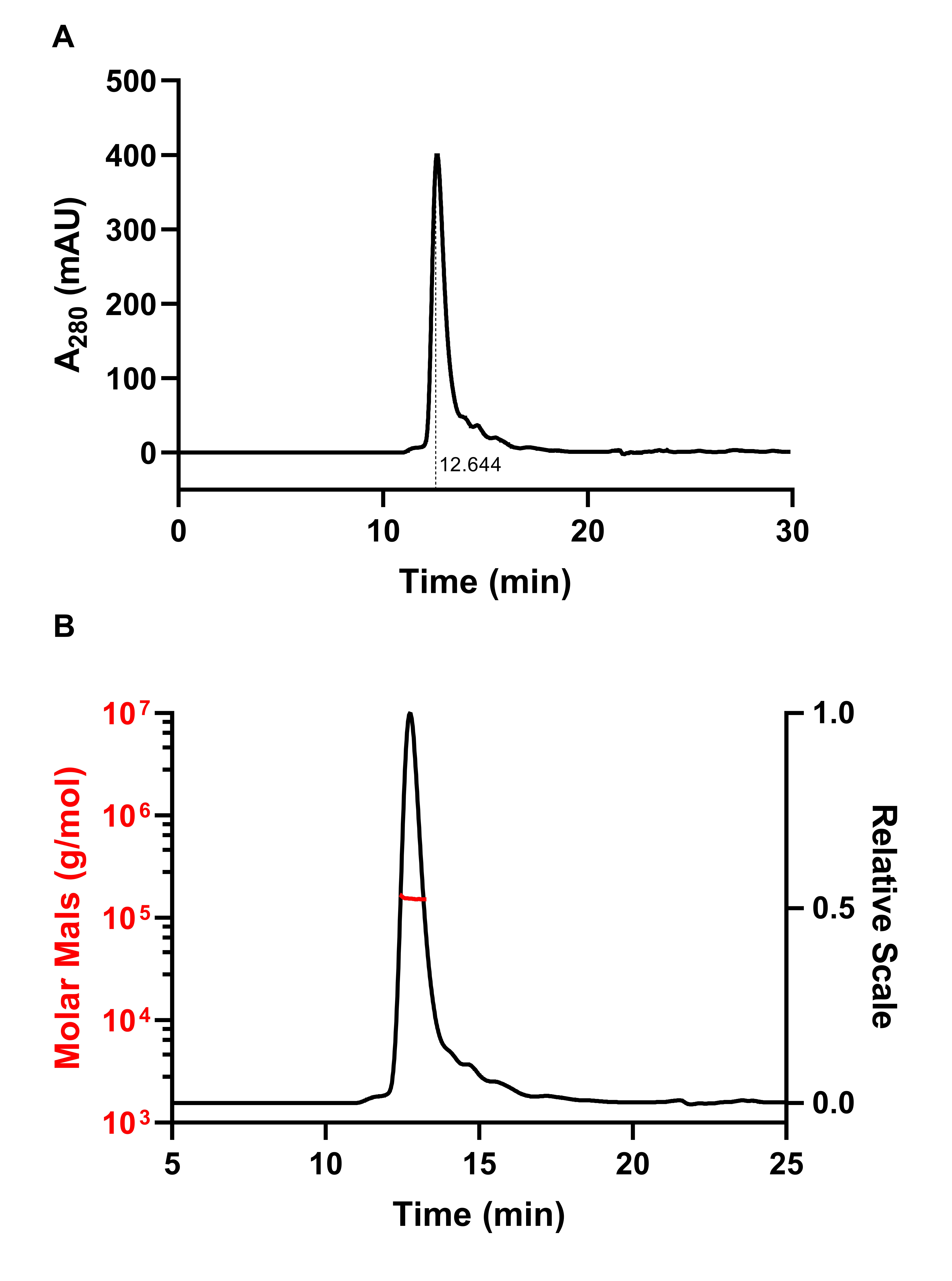

Fig.S4. Detection of EAL by SEC-MALS. (A) Absorbance at UV 280 nm. (B) Molar mass.

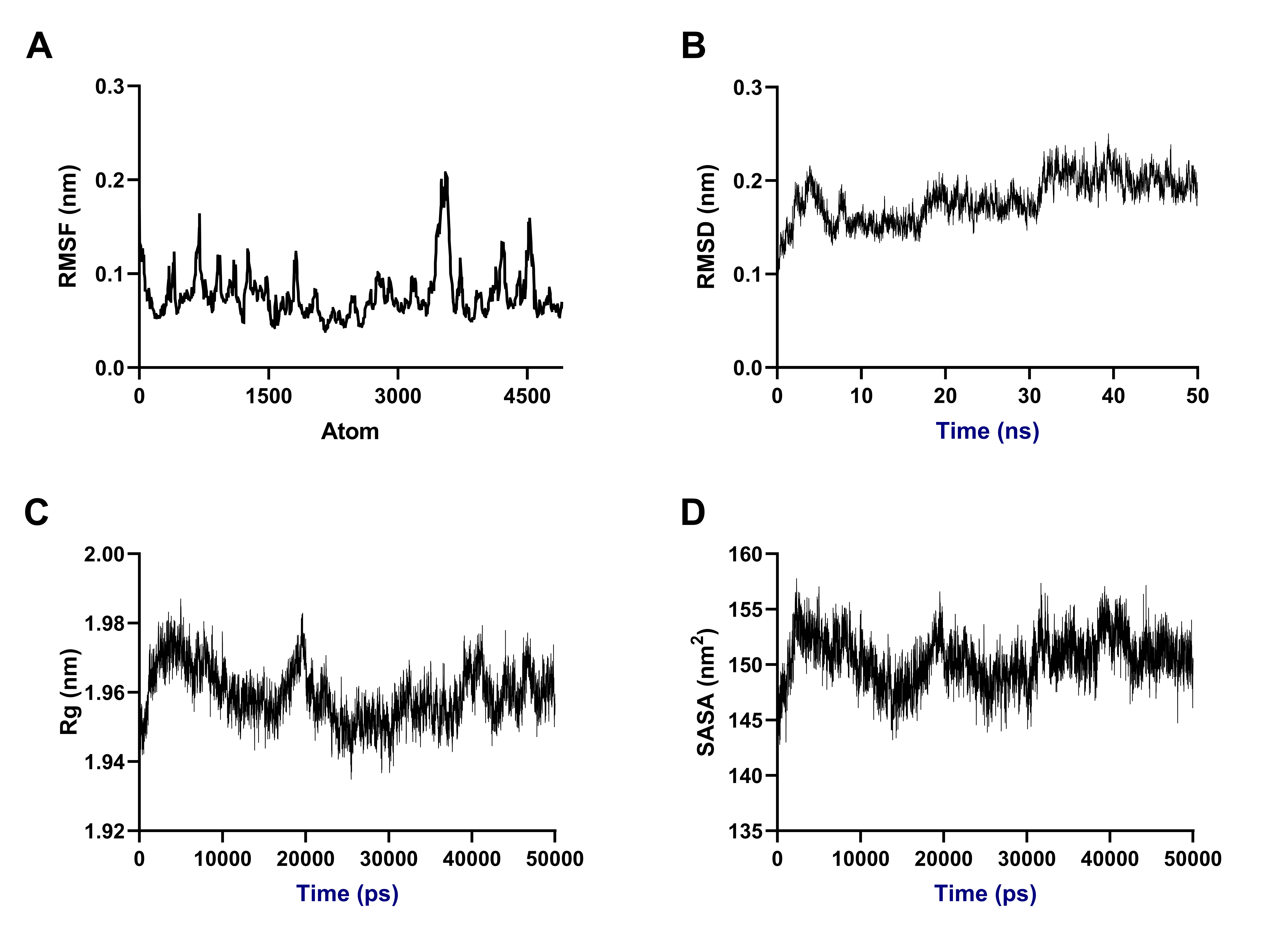

Fig. S5. Molecular dynamics simulation of EA and EAL complexes. (A) RMSF for the backbone carbon atoms of the EA and EAL complexes. (B) RMSD of the residues in the EA and EAL complexes. (C) Rg of the EA and EAL complexes. (D) SASA of the EA and EAL complexes.

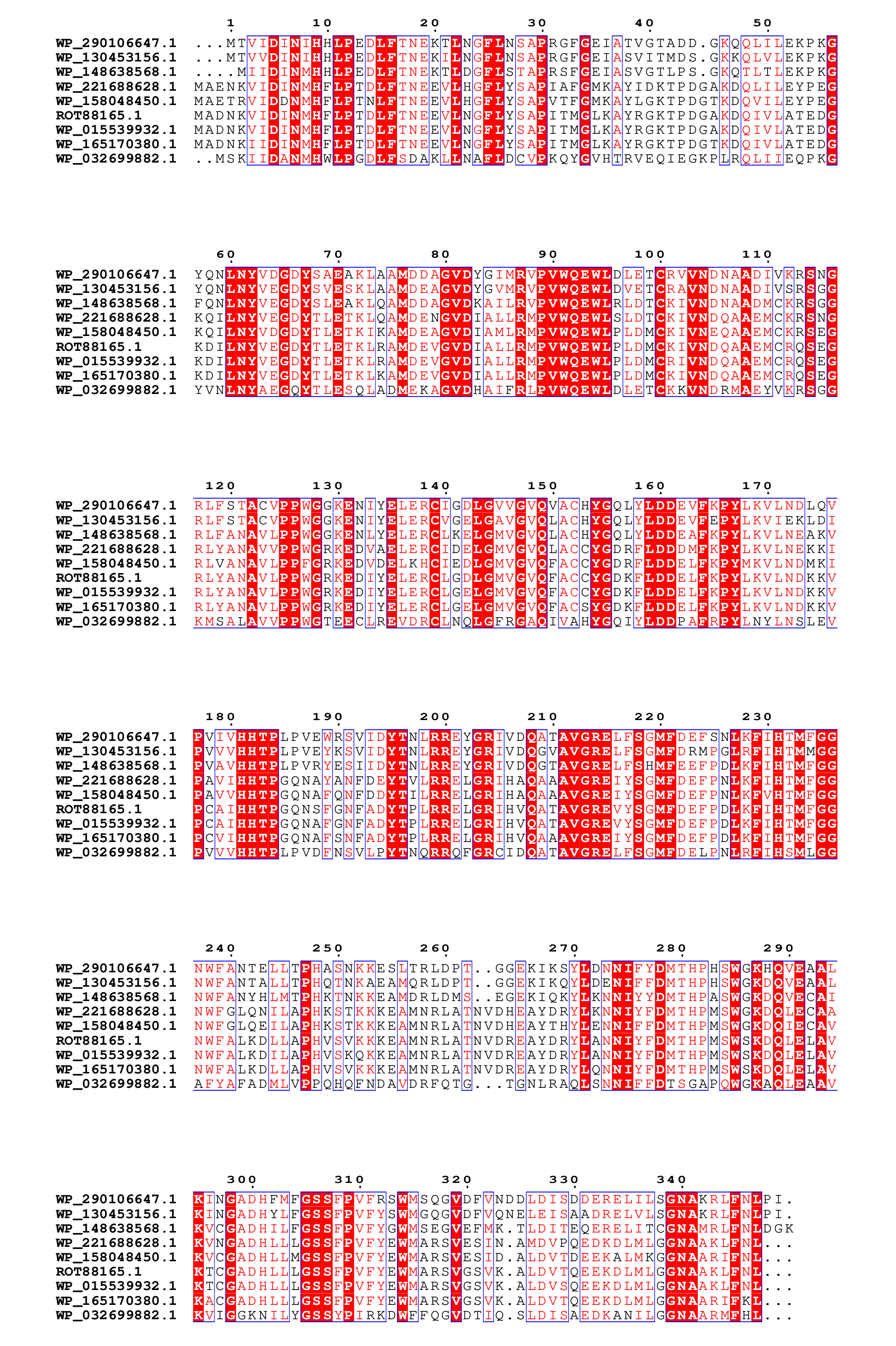

Fig. S6. Amino acid sequence alignment of EAL and validated EAL homologues was performed. The sequences include Wp_158048450.1 from *Ellagibacter isourolithinifaciens* (eis), Wp_221688628.1 from *Gordonibacter massiliensis* (gma), Wp_148638568.1 from *Acetobacterium wieringae* (awi), Wp_015539932.1 from *Gordonibacter pamelaeae* (gpa), Wp_165170380.1 from *Adlercreutzia* sp. ZJ242 (asp), Wp_032699882.1 from *Raoultella planticola* (rpl), Wp_290106647.1 from *Microbacterium oryzae* (mor), and Wp_130453156.1 from *Leucobacter luti* (llu).
